# Characterizing substructure via mixture modeling in large-scale genetic summary statistics

**DOI:** 10.1101/2024.01.29.577805

**Authors:** Hayley R Stoneman, Adelle Price, Nikole Scribner Trout, Riley Lamont, Souha Tifour, Nikita Pozdeyev, Colorado Center for Personalized Medicine, Kristy Crooks, Meng Lin, Nicholas Rafaels, Christopher R Gignoux, Katie M Marker, Audrey E Hendricks

## Abstract

Genetic summary data are broadly accessible and highly useful including for risk prediction, causal inference, fine mapping, and incorporation of external controls. However, collapsing individual-level data into groups masks intra- and inter-sample heterogeneity, leading to confounding, reduced power, and bias. Ultimately, unaccounted substructure limits summary data usability, especially for understudied or admixed populations. Here, we present *Summix2*, a comprehensive set of methods and software based on a computationally efficient mixture model to estimate and adjust for substructure in genetic summary data. In extensive simulations and application to public data, *Summix2* characterizes finer-scale population structure, identifies ascertainment bias, and identifies potential regions of selection due to local substructure deviation. *Summix2* increases the robust use of diverse publicly available summary data resulting in improved and more equitable research.

## Introduction

Genomic sumsmary data advances both basic research and clinical translation. These often take the form of allele frequencies (AFs) or Genome Wide Association Study (GWAS) test statistics, and are used to prioritize putative causal genetic variants, as external common controls^1,2^, for causal inference via Mendelian randomization^3^, and for risk prediction using polygenic scores (PGS)^4^. Compared to individual-level data, summary data is easier to access, store, and analyze while protecting individual data privacy. This enables reuse and subsequently greater opportunities for genetic discovery^1^.

However, broad, rigorous, and representative use of summary data has been historically limited by an inability to identify and account for substructure within and between datasets. Indeed, issues with the use of summary data are often most pronounced for ancestrally admixed and diverse populations. While efforts are actively underway to collect more representative samples including those from admixed populations (e.g., African American, Latino), the majority of genetic research thus far has been performed in groups of European ancestry and is not fully generalizable to diverse or admixed populations^5-9^. Further, existing admixed data are often underutilized due to barriers in completing robust research using summary data with genetic substructure^1,10^. The 2023 National Academy report on *Population Descriptors in Genetics and Genomics Research* recommends the use of genetic similarity (i.e., quantitative measures of shared genetic ancestry) to prevent confounding in most genetics and genomics research while also recognizing that estimating genetic similarity is often not possible with summary data^11^. Thus, researchers or clinicians working with admixed or underrepresented ancestral groups often face an unfair choice: choose the best (but poorly matched) public summary data and produce potentially biased results, or do not complete the study. This choice exacerbates inequities in research and, ultimately, in precision medicine and healthcare derived from the research. Further, a comprehensive understanding of fundamental biological processes is also constrained by incomplete use of diverse and representative data^12-14^.

Numerous methods exist to detect global (i.e., average across the genome) and local (i.e., windows across the genome) ancestral substructure in admixed populations using individual level data. Methods to infer global population structure include principal component analysis^15^, ADMIXTURE^16^, and identity by descent (IBD) clustering^17-19^. Local ancestry methods include LAMP-LD^20^, RFMix^21^, ELAI^22^ and Loter^23^. While both methods can be used to adjust for population structure, studies suggest that global ancestry adjustment is usually optimal to control for type I error and retain statistical power, while local ancestry adjustment is better at improving fine-mapping and accuracy of effect size estimates^24-27^. In addition, local ancestry methods have been used to identify signatures of post-admixture selection such as the HLA locus, which is known to play a role in immune response ^22,28-35^.

While many methods exist to detect and account for substructure using individual level data, few methods exist for summary level data. In 2021, we developed *Summix*, a computationally efficient mixture model employing AFs to estimate and adjust for continental level ancestry substructure in genetic summary data^36^. In 2022, Privé presented a similar method in the *bigsnpr* package that minimizes AF projections onto a principal component (PC) space derived from individual level UK Biobank data^37^, enabling identification of finer-scale ancestral genetic substructure. In contrast, *Summix* does not require individual level data and can estimate substructure with far fewer variants (e.g., ∼100) opening the possibility of additional functionalities such as estimation of local substructure and genetic similarity to case status.

Here we present *Summix2*, a comprehensive suite of methods to detect and leverage substructure in genetic summary data. Advances include estimating finer-scale and local substructure across the genome, an effective sample size for adjusted AFs, generalizing a goodness-of-fit (GoF) metric for any array or sequencing technology, estimating other types of genetically driven substructure such as genetic predisposition to conditions, and identifying regions where local substructure differs from the average global (to search for possible genomic signals of selection). *Summix2* is available as an R package on Bioconductor with open-source code and extensive examples to ensure reproducibility and enable others to use and build on our methods.

## Results

### Summix2 overview

*Summix*^*36*^ employs a mixture model to estimate substructure proportions in genetic summary data by minimizing the least-squares (LS) difference between a vector of observed sample AFs and vectors of reference group AFs. Using the estimated substructure proportions, *Summix* can adjust observed AFs to match target sample subgroup proportions, supporting harmonization of summary data. *Summix2* has significantly expanded capabilities including a GoF metric generalizable across different data generation technologies, an effective sample size estimate for adjusted AFs, estimation of local substructure with a test to identify regions where local substructure differs from the average, and generation of a finer-scale genetic similarity map for improved data harmonization (Fig. 1).

**Fig. 1:**
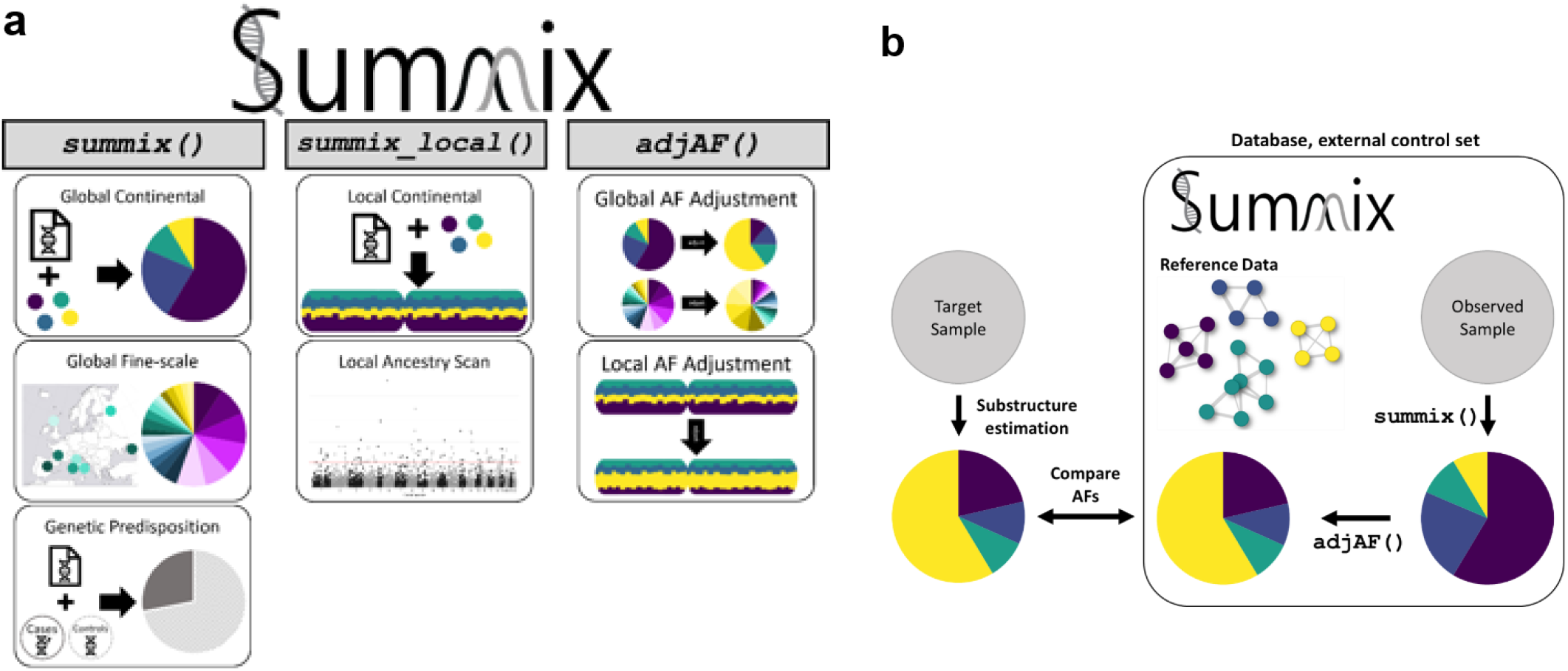
*Summix2* suite of methods: overview of functions and uses. **a**, Three functions to detect and leverage substructure in genetic summary data. The *summix()* function estimates substructure proportions given AFs for the observed and reference groups, *summix_local()* detects and leverages local substructure proportions, and *adjAF()* adjusts AFs for an observed sample to match the substructure proportions of a target sample or individual. **b**, a representation of the *Summix* workflow.

### Generalized goodness-of-fit metric

The GoF metric for the original *Summix* method was the sum of the LS objective function and was developed using gnomAD v2.1 whole-exome data^36^. We applied *Summix* to gnomAD v3.1.2 whole genome sequencing data^38^ and identified that the original GoF metric was sensitive to the AF spectrum. Through simulations, we found that while substructure estimate accuracy was stable across MAF bins, the original GoF metric increased with MAF (Extended Data Fig. 1). To remove correlation between the GoF metric and MAF, we weighted the LS loss by MAF bin (bins: (0, 0.1], (0.1, 0.3], (0.3-0.5], weights: 5, 1.5, 1, respectively). In simulations iteratively leaving out one reference group with varying substructure proportions from the reference panel, we found the new GoF metric was <0.5 when ≤10% of substructure was not accounted for by the reference panel and was >1.5 when ≥20% was not accounted for. Thus, we propose the same thresholds as the original GoF metric: <0.5 good fit, 0.5-1.5 moderate fit, and >1.5 poor fit. Additional details are in **Online Methods**.

### Experiment overview

We evaluated *Summix2* using AF data from gnomAD v3.1.2^38^ as the observed summary data. This data includes whole genomes from 76,156 individuals and ten gnomAD groups (i.e., African/African American (AFR/AFRAM), Amish (AMI), Latino/Admixed American (AMR), Ashkenazi Jewish (ASJ), East Asian (EAS), Finnish (FIN), Non-Finnish European (NFE), Middle Eastern (MID), South Asian (SAS), and Other (OTH)) (Supplementary Table 1). For reference AF data, we used 73 finer-scale reference groups from the 1000 Genomes (1KG)^39^ and Human Genome Diversity Project (HGDP)^40^ that was harmonized and released with gnomAD v3.1.2^38^. We removed 21 admixed reference groups (i.e., with notable contributions from more than one continental grouping)^41^, one group not in the five continental groups used (HGDP Kalash), and one group with N=1 (HGDP Surui) resulting in 50 finer-scale reference groups. Using gnomAD v3.1.2 grouping assignments, five continental reference AFs (i.e., African (AFR), Non-Finnish European (EUR), East Asian (EAS), Indigenous American (IAM), and South Asian (SAS)) were created using a weighted average of the finer-scale reference AFs (Supplementary Table 2).

**Table 1:** Replicated and genome-wide significant local substructure blocks in gnomAD.

| Locus | Coordinates | Sample | P-value | Potential Gene(s) of Interest* |
| --- | --- | --- | --- | --- |
| *Replicated Signals* |  |  |  |  |
| 6p21-22 | 28749843-33594193 | AFR | 6.50e-11** | *HLA* |
| 1p33 | 48977431-49882955 | AMR | 5.49e-4 | *CYP4A11* |
| 6p21-22 | 27410678-32584612 | AMR | 1.21e-5 | *HLA* |
| *Genome-Wide Scan* |  |  |  |  |
| 1q42.13*** | 226810857-227371817 | AFR<br>AMR | 2.18e-8<br>5.11e-8 | *PSEN2, COQ8A* |
| 2p11.2 | 89085547-91505258 | AFR<br>AMR | 1.19e-6<br>1.48e-6 | *IGKV* |
| 1q32.2 | 209710004-210350566 | AFR | 1.99e-6 | *LAMB3, HSD11B1, IRF6, SYT14* |
| 3p21.31 | 49230341-50009909 | AFR | 2.78e-6 | *RHOA, DAG1, TRAIP, MST1R* |
| 10q24.2 | 98531740-99215582 | AFR | 1.08e-5 | *HPSE2* |
| 11p11.2 | 48713155-49141143 | AFR | 1.80e-9 | *TRIM* |
| 18q12.2 | 39086699-39784890 | AFR | 5.79e-10 | *MIR924HG* |
| 3q22.1 | 132492441-133141527 | AMR | 2.69e-6 | *UBA5, NPHP3* |
| 5p13.1 | 41972605-42576421 | AMR | 5.32e-9 | *GHR* |
| 9p21.3 | 20543715-21637268 | AMR | 6.77e-8 | *IFNA, MIR31HG* |
| 9q33.3 | 125309605-125966124 | AMR | 7.03e-7 | *MAPKAP, PBX3* |
| 13q14.11 | 40546196-41169051 | AMR | 2.09e-6 | *FOXO1, SLC25A15* |

\* Full list of genes is in Supplementary Table 6
\*\* Replication Bonferroni-corrected significance threshold < 1.56e-3 AFR, 5.75e-4 AMR
\*\*\*Genome-wide Bonferroni-corrected significance threshold < 4.21e-6 AFR, 2.80e-6 AMR

**Table 2:** GO Enrichment Analysis for genes in genome-wide significant blocks in gnomAD groups.

| gnomAD Group | GO Biological Process | P-value | FDR | Involved in Immunity? |
| --- | --- | --- | --- | --- |
| AFR/<br>AFRAM | immunoglobulin production | 4.29E-27 | 5.18E-18 | Yes |
|  | immune response | 2.09E-23 | 8.41E-17 | Yes |
|  | meiotic DNA double-strand break formation | 1.30E-07 | 4.79E-05 | No |
|  | response to reactive oxygen species | 5.21E-06 | 1.37E-03 | Yes |
|  | innate immune response | 2.40E-04 | 5.32E-02 | Yes |
|  | positive regulation of adaptive immune response | 2.29E-03 | 4.59E-01 | Yes |
|  | antigen processing and presentation | 3.00E-03 | 5.50E-01 | Yes |
|  | protein ubiquitination | 3.50E-03 | 5.51E-01 | Yes |
|  | regulation of leukocyte mediated cytotoxicity | 3.51E-03 | 5.16E-01 | Yes |
|  | positive regulation of lymphocyte mediated immunity | 3.79E-03 | 4.90E-01 | Yes |
|  | regulation of receptor signaling pathway via JAK-STAT | 4.33E-02 | 1.00E00 | Yes |
|  | synapse assembly | 4.33E-02 | 1.00E00 | No |
| AMR | immunoglobulin production | 2.17E-23 | 4.79E-20 | Yes |
|  | immune response | 2.44E-15 | 1.79E-12 | Yes |
|  | response to stimulus | 6.18E-04 | 1.67E-01 | Yes |
|  | neuron projection extension | 2.04E-02 | 1.00E00 | No |
|  | ubiquinone biosynthetic process | 2.29E-02 | 1.00E00 | Yes |
|  | positive regulation of receptor signaling pathway via JAK-STAT | 2.29E-02 | 1.00E00 | Yes |
|  | postsynapse organization | 2.80E-02 | 1.00E00 | No |
|  | regulation of cytokinesis | 4.29E-02 | 1.00E00 | No |

PANTHER GO-Slim Biological Process annotations and p-values for the set of genes in all genome-wide significant blocks from the local substructure scan in gnomAD AMR and AFR/AFRAM. Only processes with $P < 0.05$ are included in this table.

We used simulated observed and reference AF data to assess the accuracy and precision of substructure estimates, absolute and relative difference of AF adjustments, and type I error and power to detect regions where local substructure differs from global. Simulated AF data was generated using a multinomial distribution with reference AFs as the input probabilities. Observed admixed data was generated using mixing proportions of the simulated reference group AFs with a default sample size of 10,000. Non-admixed reference data was simulated from one reference group with a default sample size of 500 individuals for continental references and 100 for finer-scale references.

For local substructure simulations, linkage disequilibrium (LD)-filtered (r^2^<0.2) chromosome 19 variants and 100 simulation replicates were used for each simulation scenario. Simulation parameters included varying mixing proportions, level of substructure (i.e., continental vs. finer-scale), reference sample size, observed sample size, region size with differing substructure proportions, and window size. For power to detect local substructure, we simulated a 50Kb to 1Mb region with different mixing proportions. We used the remainder of the chromosome to evaluate type I error.

We estimated the accuracy and precision of finer-scale substructure detection for reference groups with varying degrees of similarity (as estimated by the Hudson estimator for pairwise *F*_*ST*_)^42^ using a random subset of 100,000 genome-wide variants and 100 simulation replicates. Local and finer-scale simulations result in many substructure estimates (e.g., windows, reference groups). We report the *minimum accuracy*, which we define as the minimum accuracy across the set of estimates (e.g., across all windows and reference groups).

Full details are in **Online Methods**.

### Overview of Summix-local method

By scanning regions of the genome, *Summix* can be used to detect local substructure as well as regions where the local substructure differs from global. Within the *Summix-local* method, we employ two algorithms for scanning the genome: a classic sliding windows approach, and a fast catch-up algorithm that enables dynamic window sizes^43^. To identify regions where local substructure differs from the average substructure across the chromosome (like in admixture mapping^44^), we developed a Wald test statistic. Standard error was estimated as an average of simulated within block standard error and observed between block standard deviation. See **Online Methods** for details.

### Summix-local produces accurate and precise local substructure estimates

*Summix-local* produced local substructure estimates with high accuracy and precision in simulated AMR-like and AFR/AFRAM-like admixed samples. The minimum accuracy across blocks was high (>99.5%) for simulation scenarios with moderate to large reference sample sizes (N≥500), all observed sample sizes evaluated (N≥1000), and all window sizes evaluated (≥250 variants). For simulations with small reference sample sizes (N=100 per group), the minimum accuracy decreased slightly (98%) (Fig. 2a). Minimum precision for all scenarios was high (>98%); and any reduction in precision was seen for smaller sample and window sizes (Supplementary Fig. 1). We assessed the impact of simulated LD on local substructure estimates (as described in Supplementary Note 1, Supplementary Fig. 2). LD did not substantially impact accuracy but did result in a lower number of effectively independent single nucleotide polymorphisms (SNPs) per window, artificially reducing the standard error, which could lead to inflated test statistics and false positives. Therefore, we recommend using LD filtered data for *Summix-local*.

**Fig. 2:**
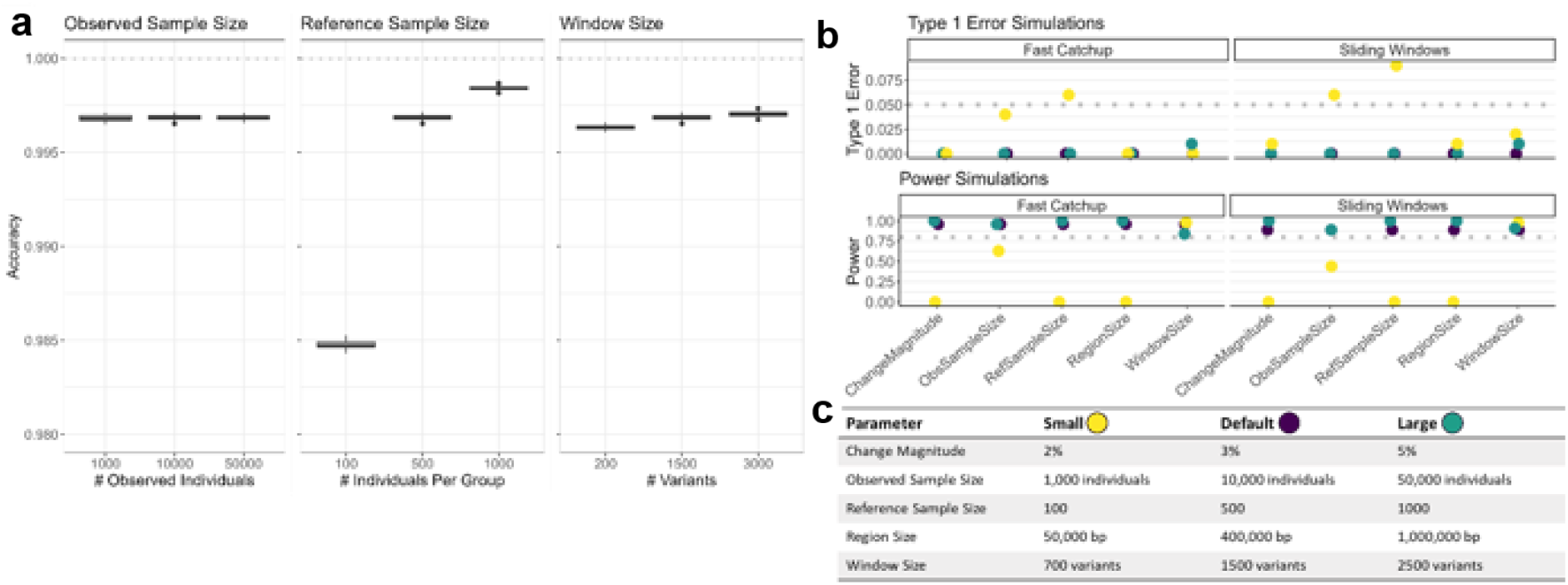
*Summix-local* accurately estimates local substructure proportions and detects regions where local substructure differs from global in an AMR-like simulation. **a**, Accuracy (y-axis) for *Summix-local* for 100 replicates per scenario using default simulation parameters (1000 observed individuals, 500 individuals per reference group, window size 1500 variants) except when varying by: observed sample size (left), reference sample size (middle), and window size defined by number of variants (right). **b**, Type I error (top) and power (bottom) for the *Summix-local* substructure scan for fast-catchup (left) and sliding windows (right) algorithms across simulation scenarios as shown in **c** using LD filtered (r^2^<0.2) chromosome 19 data. For **b** and **c** the parameter settings are default except where the given parameter is changed to be small or large.

### Ability to detect regions with local substructure deviations

We then assessed the type I error and power of *Summix-local* to identify regions of the genome where local substructure deviates from the chromosome average. The type I error rate was conservative, below the expected 5% rate, for most simulation scenarios but approached or exceeded 5% for small reference or observed sample sizes (N=100 and N=1000, respectively) in the AMR-like sample (AMR-like: Fig. 2b,c, Supplementary Table 3; AFR/AFRAM-like: Extended Data Fig. 2, Supplementary Table 4). *Summix-local* maintained high power (>80%) given sufficiently large sample sizes (observed N≥10000, reference N≥500), regions (≥ 400kb), and magnitude of difference (≥3%). Smaller reference sample sizes (N=100), regions (50Kb), or signals (2% difference in local substructure) reduced power to near zero. Power was more robust to smaller observed sample sizes (N=1000), staying above 60%. While both scanning algorithms had well controlled type I error and high power for most simulation scenarios, the fast catch-up algorithm had consistently lower type I error and higher power compared to the sliding window algorithm. Additionally, the fast catch-up algorithm more precisely detected the true break points (Supplementary Note 2). Thus, the fast catch-up scanning algorithm is the default method for *Summix-local*. Finally, we found that the window size relative to the size of the simulated differential region affected the type I error and power. A window size of half to double the size of the region of interest maintained the highest power and lowest type I error (Supplementary Note 3, Supplementary Fig. 3). While a user can specify a window size based on their hypothesis of region size, we recommend a default window size of 400Kb as this was the average size of previously identified regions of selection. In the whole-genome sequencing data employed here, 400Kb equated to ∼1500 variants although this relationship will depend on the density of the data used.

### Local-substructure scan in gnomAD

Using *Summix-local* with the fast catch-up algorithm and default parameters (**Online Methods**), we scanned gnomAD v3.1.2 AFR/AFRAM and AMR groups for regions where local substructure differed from global substructure. In AMR, we removed four blocks with low-quality substructure estimation (i.e. GoF > 1.5) from subsequent analyses. First, we assessed regions of selection previously identified in AFR/AFRAM or AMR samples (Supplementary Table 5) ^22,29,31,32,45-48^, using a Bonferroni-corrected significance threshold for 16 and 29 regions and two or three reference groups, respectively (Fig. 3, Table 1). In both groups we replicated the *HLA* locus on chromosome 6 (6p21-23) (AFR/AFRAM *P* = 6.50e-11; AMR *P* = 1.21e-5). The *HLA* gene cluster is widely known to have functional properties under selection in both ancient and recent human populations^49^. Additionally, in AMR, we replicated a signal on chromosome 1 (1p33) containing *CYP4A11* (*P* = 5.49e-4), which is a member of the cytochrome P450 superfamily of enzymes involved in xenobiotic compound metabolism (including drugs)^31^.

**Fig. 3:**
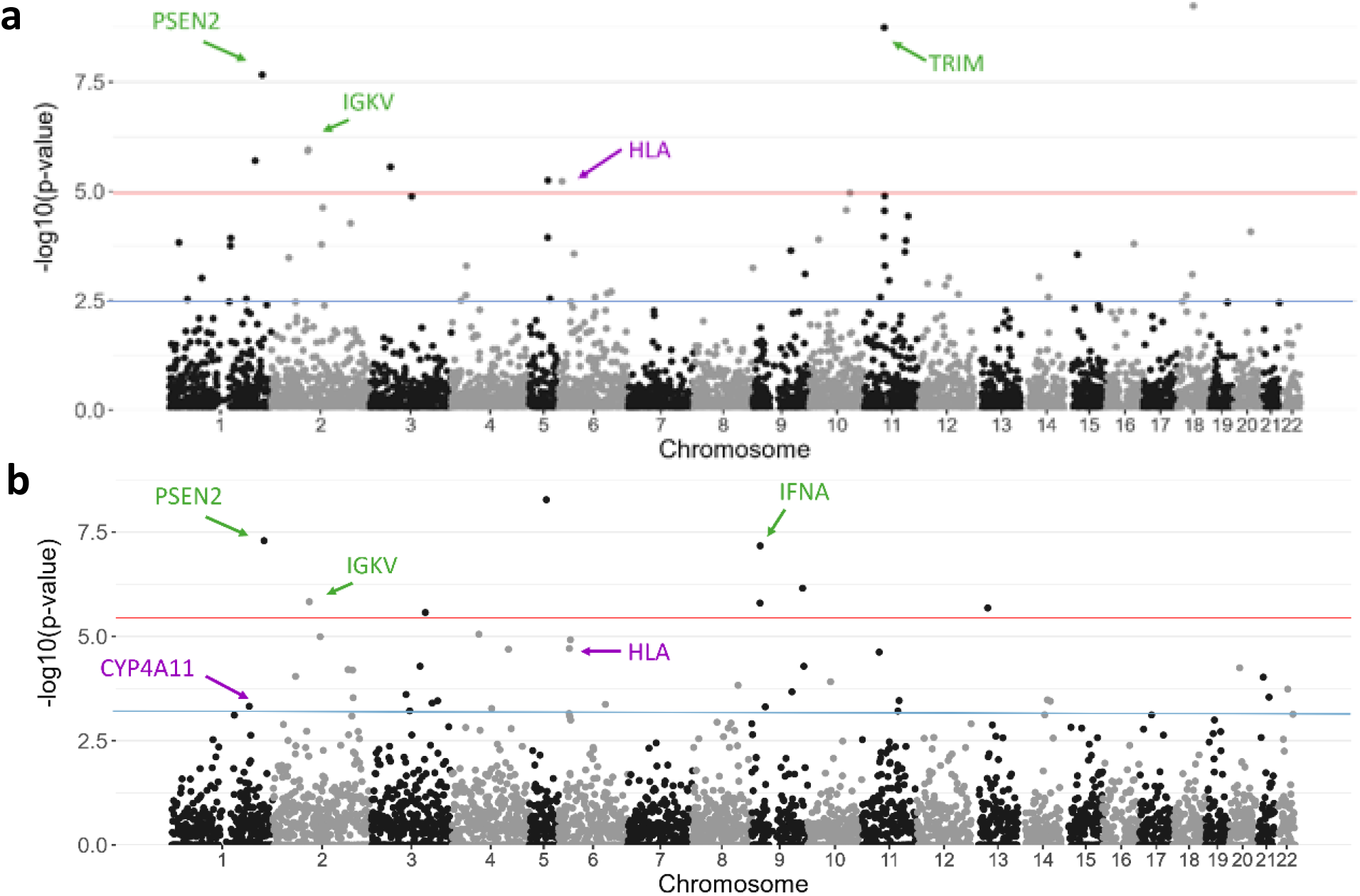
Local substructure scan in gnomAD African/African American and Latinx/Admixed American identifies potential regions of selection. Fig. 3: Local substructure scan in gnomAD African/African American and Latino/Admixed American identifies potential regions of selection. *Summix-local* was applied to gnomAD v3.1.2 using the fast-catchup algorithm and default parameters. **a**, African/African American (AFR/AFRAM) and **b**, Latino/Admixed American (AMR). Red line indicates Bonferroni-corrected genome-wide significance (*P* < 4.21e-6 AFR, 2.80e-6 AMR); blue indicates Bonferroni-correction for the replication signals (*P* < 1.56e-3 AFR, 5.75e-4 AMR). Genes of interest discussed in the **Results** are annotated; for a complete list of genes see Supplementary Table 6. Gene names in purple are replicated signals that were previously identified, while green are putative novel signals.

In a genome-wide scan, we detected nine and eight significant blocks representing eight and seven putative regions of selection in the AFR/AFRAM and AMR groups, respectively. Genome-wide significance was defined by a Bonferroni-correction for 5945 blocks and two or three reference groups resulting in thresholds of 4.21e-6 and 2.80e-6, respectively for AFR/AFRAM and AMR (Table 1, Supplementary Table 6). Two of the loci, 1q42.13 and 2p11.2 (AFR/AFRAM *P* = 2.18e-8, 1.19e-6; AMR *P* = 5.11e-8, 1.48e-6, respectively), were identified in both groups. The locus at 1q42.13 contains 12 genes including *PSEN2*, which has been associated with Alzheimer’s disease^50^. The locus at 2p11.2 contains 32 total genes, including the immunoglobulin kappa variable cluster (*IGKV*), which is involved in immune response and has been associated with breast cancer^51^. The 9p21.3 locus was identified in AMR (*P* = 6.77e-8) and contains 41 genes including a cluster of alpha interferon genes, *IFNA* and *MIR31HG*. These alpha interferons are a key part of the innate immune response. *MIR31HG* encodes a long non-coding RNA that has been associated with cancers including pancreatic, lung, and bladder, as well as autoimmune disease^52-55^. The 11p11.2 locus was identified in AFR/AFRAM (*P* = 1.80e-9) and contains a block of *TRIM* genes among a total 20 genes, which are predicted to be involved in protein and ion binding and the innate immune response. SNPs in these genes have been associated with glomerular filtration rate, chronic kidney disease, and BMI^56,57^. Using Gene Ontology (GO) Enrichment Analysis, we found enrichment for genes in immune response pathways including immunoglobulin production (AFR/AFRAM FDR = 5.18e-18, AMR FDR = 4.79e-20) and immune response (AFR/AFRAM FDR = 8.41e-17, AMR FDR = 1.79e-12) (Table 2)^58,59^.

### Detecting finer-scale genetic substructure

We estimated genetic similarity for the 50 finer-scale and five continental reference groups using the Hudson estimator^42^ for pairwise *F*_*ST*_ (Extended Data Fig. 3). Pairwise *F*_*ST*_ estimates were generally greater, indicating less similarity, between continental groups (ranging from 0.026 to 1.31) compared to finer-scale groups (0.001 to 0.035). Exceptions included higher similarity between EUR and SAS continental groups (0.026) and less similarity between finer-scale reference pairs such as Colombian and Pima (0.07) within the IAM continental reference grouping (Supplementary Tables 7 and 8).

We evaluated the ability of *Summix2* to estimate substructure proportions for increasingly similar pairs of finer-scale groups, i.e., decreasing pairwise *F*_*ST*_. We simulated three mixture proportion scenarios (i.e., four reference groups simulated at equal, varying, and one zero with three equal mixing proportions) across three pairwise *F*_*ST*_ scenarios (pairwise *F*_*ST*_= 0.009, 0.007, 0.005) using EAS finer-scale reference groups. We estimated substructure proportions using the 50 finer-scale reference panel. Minimum accuracy was inversely related to similarity; accuracy decreased as finer-scale group similarity increased (i.e., *F*_*ST*_ decreased, Fig. 4a). The minimum precision of *Summix2* estimates was consistently high (>99%) (Supplementary Table 9). Importantly, in all simulation scenarios, highly similar finer-scale reference groups captured the bias in the mixture proportion estimates. The genetic similarity map reflects this result, showing the bias in the reference group proportion estimates is distributed locally within the network to other similar nodes (i.e., reference groups) (AFR/AFRAM-like, Fig. 4b,c; EAS-like, Extended Data Fig. 4).

**Fig. 4:**
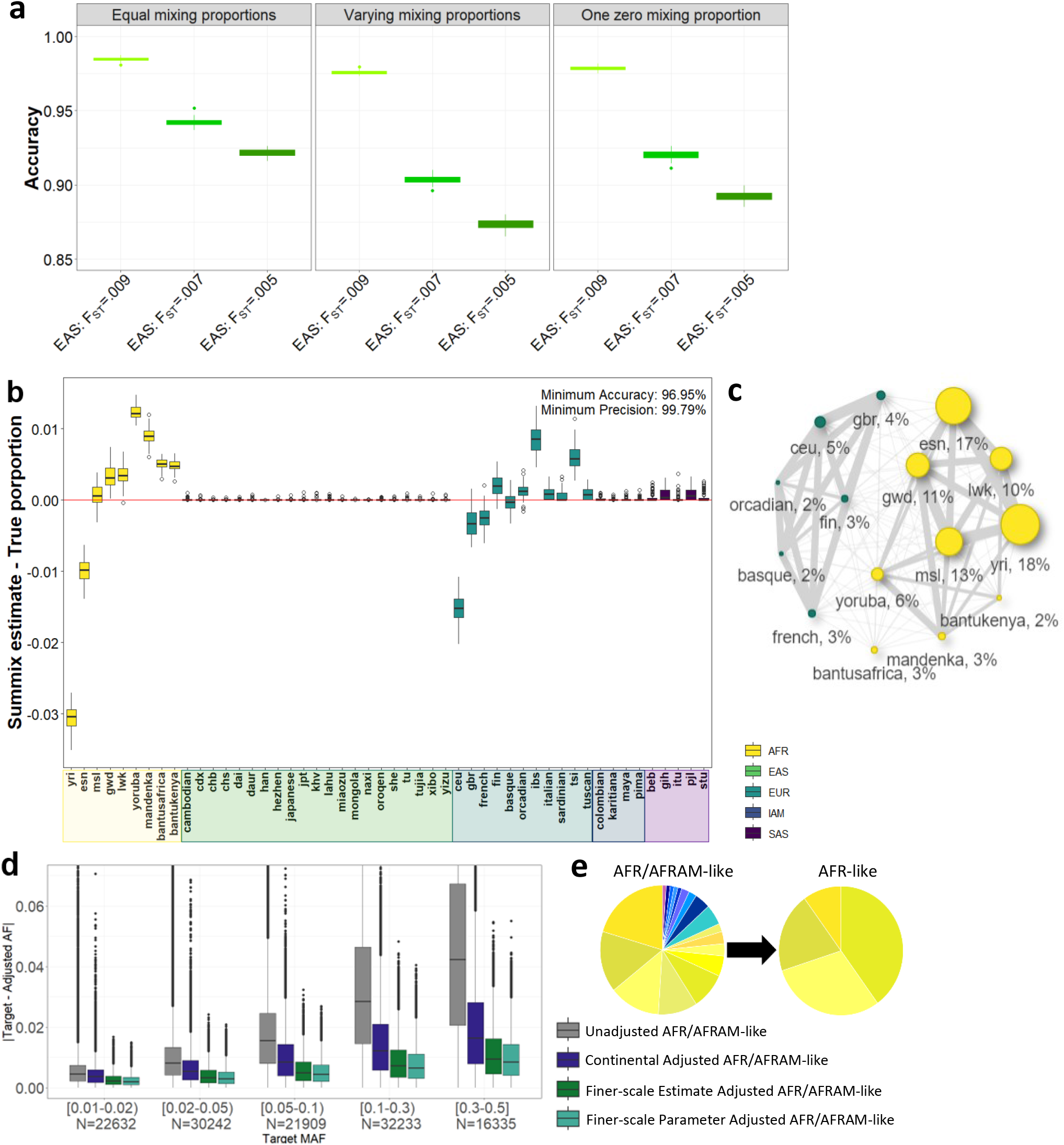
*Summix2* detects and adjusts for finer-scale genetic substructure. **a**, Accuracy of *Summix2* estimates for three simulation scenarios: Equal Proportions (.25, .25, .25, and .25), Varying Proportions (.1, .2, .3, and .4), and One Zero proportion (0, .33, .33, .33) across pairwise groups of increasing similarity (***F***_***ST***_ =0.009, light green, low similarity; ***F***_***ST***_ =0.007, green, medium similarity; ***F***_***ST***_ =0.005, dark green, high similarity). **b**, Accuracy of AFR/AFRAM-like simulation scenario. **c**, Genetic substructure similarity map of *Summix2* estimates for the AFR/AFRAM-like sample where edge thickness between nodes indicates pairwise similarity (thicker edges indicate higher similarity) as defined by pairwise ***F***_***ST***_ and node size indicates magnitude of the Summix2 mixture proportion estimate for the given reference group. **d**, Absolute difference between target AFR-like sample AFs and unadjusted (gray) or adjusted observed AFR/AFRAM-like AFs grouped by MAF bin (continental adjustment, blue; finer-scale estimate adjustment, green; finer-scale parameter adjustment, teal). **e**, Finer-scale substructure proportions for AFR/AFRAM-like sample and AFR-like sample in (**d**); simulated proportions in Supplementary Table 11. (Note: “-like” nomenclature omitted from x-axis plot labels for simplicity from (**b**) and (**c**) Fig.s.)

As a resource to the community, we provide finer-scale substructure proportion estimates for seven gnomAD v3.1.2 groups with good or moderate GoF (Supplementary Table 10). Amish, Ashkenazi Jewish, Middle Eastern substructure estimates were excluded due to poor GoF (90% Confidence Interval (CI)) = 6.34 (6.16, 6.55), 2.15 (2.07,2.24), and 2.39 (2.32, 2.46), respectively).

### Adjusting for finer-scale genetic substructure using genetic similarity mapping

We next investigated adjusting the observed sample AFs so that the finer-scale substructure proportion estimates match a target sample as might be done when harmonizing or adjusting for population structure in summary datasets. We simulated an AFR/AFRAM-like observed sample and evaluated bias between the substructure proportion estimates and true simulation parameters (Fig. 4b). We then adjusted the observed AFs so that the substructure group proportions matched a simulated target AFR-like sample (simulated substructure in Supplementary Table 11). We performed the AF adjustments using finer-scale substructure estimates, continental substructure estimates, and simulated substructure proportions within the observed sample. Though the finer-scale substructure proportion estimates were biased (Fig. 4b), the AF adjustment using finer-scale substructure had lower absolute and relative difference compared to the AF adjustment using continental substructure (Fig. 4d, Extended Data Fig. 5a). Notably, the absolute difference in adjusted AFs using the biased finer-scale estimates was similar to that using the oracle simulation parameters, indicating the bias in the substructure estimates was not propagated to the AF adjustment. This is likely due to the distribution of the bias to other highly similar reference groups, as demonstrated in the genetic similarity map (Fig. 4b,c). Similar results were observed when adjusting an AMR-like sample to IAM-like using finer-scale references (Extended Data Figs. 5b-6, simulated substructure in Supplementary Table 11).

### Improved data harmonization through adjusted allele frequencies

To demonstrate the utility of *Summix2* for data harmonization under varying levels of substructure, we adjusted gnomAD AMR AFs to match the substructure for three target samples: 1KG Peruvian(Pel), a simulated target with continental substructure and Pel-like proportions (10% AFR-like, 15% EUR-like, 75% IAM-like), and a simulated target with finer-scale substructure and Pel-like proportions (45% HGDP-Maya-like, 10% HGDP-Karitiana-like, 10% HGDP-Pima-like, 10% HGDP-Colombian-like, 5% 1KG-Esan(Esn)-like, 5% 1KG-Yoruba(Yri)-like, 8% 1KG-British(Gbr)-like, 7% 1KG-Iberian(Ibs)-like) (**Online Methods**). We compared the unadjusted AFs, four *Summix2* AF adjustments (global continental, global finer-scale, local continental, and local finer-scale), and two adjustments using global and local substructure proportions released in gnomAD v3.1 that were calculated with individual level data (gnomAD global, gnomAD local. Of the 234,633 chromosome 19 variants with 1KG Pel MAF>0.01, 12,513 had local substructure estimates from gnomAD. We used this subset of variants for our comparison with gnomAD (Fig. 5) and the full set of variants for a more complete comparison of *Summix2* adjustments (Extended Data Fig. 7).

**Fig. 5:**
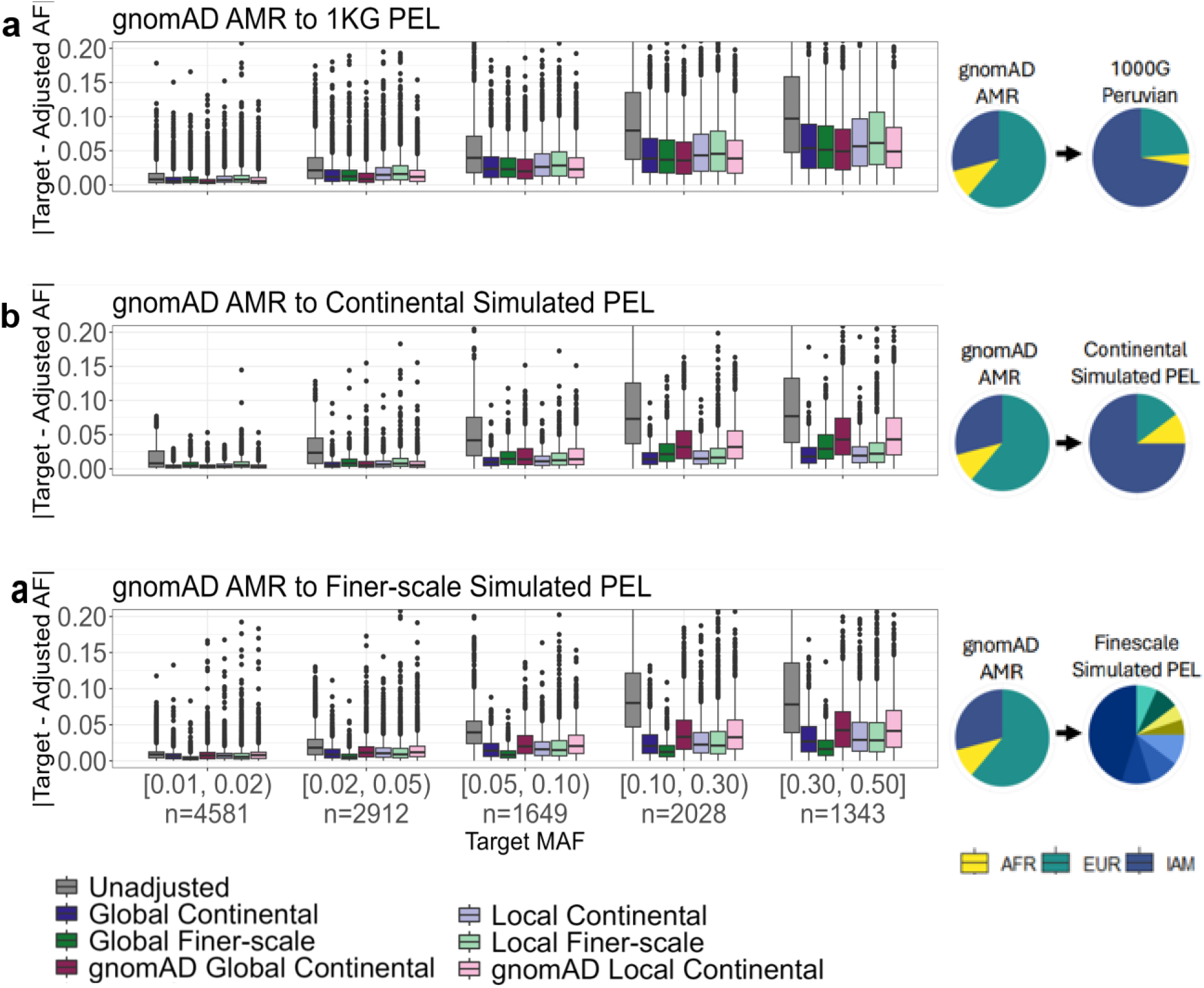
Allele frequency adjustment improves data harmonization. Absolute difference after adjusting gnomAD AMR to three different target samples **a**, 1000 Genomes Peruvian. **b**, Simulated Peruvian-like with continental-level substructure. **c**, Simulated Peruvian-like with fine-scale substructure, by target MAF bin. Results for the 12,513 variants on chromosome 19 with gnomAD local substructure estimates, and filtering for LD (r^2^ > 0.2) and MAF (>0.01). The unadjusted gnomAD AMR (grey) is compared to adjustments using gnomAD (global continental, dark red; local continental, pink) and *Summix2* (global continental, blue; global finer-scale, green; local continental, light blue; local finer-scale, light green). The global substructure proportions of the observed and target data is shown in the pie charts. The GoF was substantially better (i.e., smaller) for the simulated targets (continental substructure: continental references GoF=0.100, finer-scale references GoF=0.673; finer-scale substructure: continental GoF=1.223, finer-scale GoF=0.188) compared to the real 1KG PEL data (continental references GoF=5.488, finer-scale references GoF=5.0798), likely due to having a fully representative reference panel for the simulated targets.

All adjusted AFs had lower absolute (Fig. 5) and relative (Extended Data Fig. 8) difference compared to the unadjusted AFs (Supplementary Table 12, paired t-test adjusted *P <* 2.2e-16). We observed a larger reduction in differences when adjusting to the simulated target data compared to adjusting to the real 1KG Pel data. The reference panel is fully representative for the simulated target adjustments, likely contributing to this reduction as indicated by better GoF estimates for the simulated target data compared to the 1KG Pel data. Likely due to the large number of variants (N>12,000) the difference between most of the AF adjustments were also significantly different from each other although the differences were an order of magnitude smaller compared to improvement over the unadjusted AFs (Supplementary Fig. 4, 5). When the underlying substructure was simulated from continental-level groups, adjusting using continental reference groups, compared to finer-scale reference groups, had the lowest absolute and relative difference and highest concordance. Similarly, when the underlying substructure was simulated from finer-scale groups, adjusting using finer-scale reference groups, compared to continental reference groups, performed best (Fig. 5, Supplementary Fig. 4, 5).

The local and global adjustments generally performed similarly. One exception was higher variability and higher absolute and relative difference using local finer-scale AF adjustment compared to using global finer-scale adjustment for the simulated target with finer-scale substructure (Fig. 5, Supplementary Fig. 4, 5). The combination of a smaller number of variants and smaller reference sample sizes likely contributed to this result. Despite Summix2 using only summary data, Summix2 and gnomAD generally had similar improvement. GnomAD adjusted AFs had smaller absolute differences (<1%) and better concordance as measured by Lin’s Concordance Correlation Coefficient (CCC)^60^ (e.g., CCC = 0.9561 for gnomAD local vs CCC = 0.9400 for *Summix2* local continental) when harmonizing to 1KG Pel, but both improved the unadjusted (CCC = 0.8424) (Supplementary Fig. 6). While Summix2 outperformed gnomAD for the simulated targets, again potentially due to the same reference data being used to simulate all samples.

Finally, as the adjusted AFs are a weighted average of the observed and reference data, the sample size associated with the adjusted AFs will be different from the observed sample size. As such, we provide an effective sample size for possible use in downstream analyses such as using the adjusted AFs as external common controls in case-control association testing (**Online Methods** and Supplementary Note 4). The effective sample size for the adjusted AFs for gnomAD AMR (n=7647) ranged from 3700 to 4952 depending on the target group and reference data used (Supplementary Table 13).

### Estimating genetic similarity to case-status

We investigated using *Summix2* to detect genetic similarity to cases with a condition or disease (genetic substructure beyond ancestry). Of note, this analysis detects the conditional probability of having genetic similarity to cases given subgroup/condition status (e.g., prostate cancer cases, females). This probability is different from heritability, which estimates the probability of having a condition given a genetic background. We used case and control AFs for 138 variants from a 2018 prostate cancer PGS (**Online Methods**, Supplementary Note 5) as reference AFs to estimate the proportion of individuals in the Colorado Center for Personalized Medicine (CCPM) Biobank data that are genetically similar to the prostate cancer cases^61,62^. To match the European ancestry of the published PGS, we limited CCPM data to European-ancestry individuals, and subset by biological sex (N_female_=16,594, N_male_=10,092), prostate cancer diagnosis, and age group (i.e., ≤40 years, 41-60 years, ≥61 years) (Fig. 6). Of the males, 1317 (13%) had clinical prostate cancer diagnoses, as defined by ICD-9 and ICD-10 codes. CCPM females, who do not have prostates, can still carry genetic risk loci for prostate cancer. For CCPM females, with 0% observed prostate cancer diagnoses, estimates for the proportion genetically similar to cases ranged between 4.4% to 11.6% and did not have a trend by age group. In CCPM males with prostate cancer aged 41-60, we estimated 100% of the sample was genetically similar to reference cases (compared to reference controls). In males ≥61 with prostate cancer, the estimate was 86.5%, potentially indicating increased environmental contribution to prostate cancer risk in this group. Interestingly, for all males, the estimated proportion genetically similar to cases increased with age, from 1.8% for ≤40 years to 7.2% for 40-60 years, and 15.8% for ≥61 years, mirroring the increase in the observed prostate cancer diagnosis proportions as age increased. Since the proportion of people with a genetic predisposition to disease is not expected to increase with age (as evidenced by the observation in the female group), the increase in the proportion of males with a genetic predisposition to prostate cancer by age group likely reflects ascertainment bias, where older males were more likely to have prostate cancer and subsequently came to the University of Colorado for healthcare, where they were recruited into the CCPM Biobank. To confirm the ascertainment theory, we evaluated the proportion of non-prostate cancer males that were genetically similar to prostate cancer cases finding 1.94% for males ages ≤40 years, 4.32% for males ages 40-60 years, and 0% for males ages ≥61 years. This indicates a small proportion of unrealized genetic prostate cancer risk in non-prostate cancer males ages ≤60 further but not for the >60 group supporting that the increase in genetic similarity to prostate cancer cases by age in the all male sample is due to increased ascertainment of prostate cancer cases by age.

**Fig. 6:**
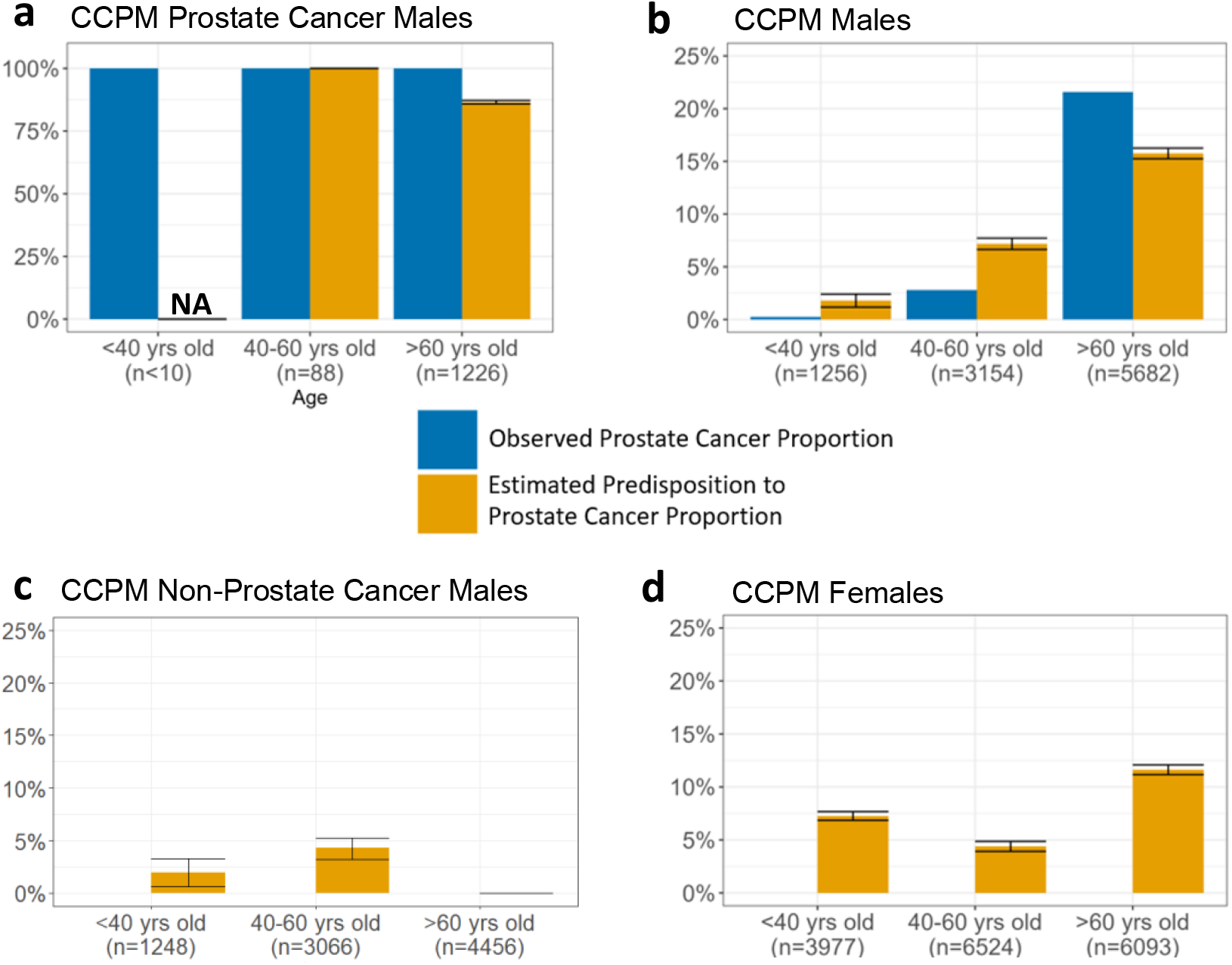
Detecting genetic similarity to prostate cancer cases in CCPM Biobank groups. Genetic substructure estimates from *Summix2* in CCPM Biobank groups using prostate cancer cases and controls as reference data for 138 variants from a 2018 prostate cancer PGS stratified by age (x-axis) and **a**, Males with prostate cancer (n = 1,322), **b**, All males (n = 10,092), **c**, Non-prostate cancer males (n = 8,770) and **d**, Females (n = 16,594). The observed proportion of individuals with prostate cancer (blue) is compared to the estimated proportion genetically similar to the prostate cancer cases reference data (orange) with jackknife 95% confidence intervals shown. There were too few males with prostate cancer under the age of 40 years (N<10) to estimate substructure proportions.

## Discussion

Here, we present *Summix2*, a Bioconductor^63^ software package and suite of methods to estimate and adjust for substructure in summary level genetic data. *Summix2* enables estimation and testing of local substructure, estimation of finer-scale genetic substructure, genetic similarity mapping, and harmonization of AFs between samples. Using prostate cancer as an example, we show that *Summix2* can also detect genetically driven substructure beyond ancestry. These tools can be used to improve genetic summary data harmonization and increase equitable use of genetic summary data across diverse groups.

Applying *Summix-local* to gnomAD AFR/AFRAM and AMR groups, we identified eight and seven putative regions of selection in a genome-wide scan, respectively, of which one and two replicated previously-identified regions of selection^22,29,32,45^. Five out of eight regions in AFR/AFRAM and three out of seven regions in AMR contain genes involved in immune response representing enrichment in the immune response pathway. Other previously identified regions of selection may have failed to replicate due to several reasons including differences in selection pressures or underlying ancestral populations. For instance, the gnomAD AMR and AFR/AFRAM groups are each an aggregation of many highly diverse samples. Specifically, samples from Mexican, Costa Rican, Colombian, Karitiana, Maya, Puerto Rican, and Peruvian populations are included in the gnomAD AMR group. These groups have different histories, admixture events, and environments likely resulting in different exposures and selective pressures. This also highlights the inadequacy of using a single group label, which falsely suggests that all subgroups share similar genetic, social, and environmental backgrounds^11^. Subsequently, the signals found in gnomAD AMR using *Summix-local* are likely strong and widely observed across most of the AMR sub-groups. The *HLA* region, identified in both the AFR/AFRAM and AMR groups, exemplifies one such robustly observed region of selection^49^. The enrichment of genes involved in the immune response pathway may also reinforce the evidence of regions of selection^58,59^. We do acknowledge that local ancestry-based methods can have additional biases, such as inconsistency in ancestry information across the genome leading to bias in local substructure estimation, however we expect our GoF filtering to help identify regions with noisy estimates. Nonetheless we describe these regions as suggestive of selection.

Although researchers have commonly used continental ancestral groupings to detect and control for population structure, genetic ancestry is not discrete and the use of groups has the potential to exacerbate existing inequities^15,16,24-27^. Further, the use of group labels paired with inconsistent use of race, ethnicity, and ancestry terms in genetics research can perpetuate the harmful belief that racial differences reflect biological differences^11,64^. The 2023 National Academies Report^11^ recommends using genetic similarity, rather than discrete groups, to control for population structure. While easily achievable with individual level data (e.g., using principal components or variance components)^15,16^, methods to model genetic similarity within summary level data are not commonly available. By creating a genetic similarity map of finer-scale reference groups in summary data and using this map to adjust and control for population structure, *Summix2* offers a step towards modeling and adjusting for genetic similarity in summary data.

Though the mapping of genetic similarity is dependent on reference data, we find that estimating genetic ancestry is more reliant on representation in the entire map rather than individual reference groups. For instance, while there is bias in subgroup proportion estimation of individual groups when using highly similar, finer-scale reference groups, the bias reflects shifts in substructure estimates to similar reference groups in the network thus creating a more continuous representation of the genetic space. Further, this bias does not propagate to AF adjustments and harmonization (at least for MAF>1% as evaluated here). More research is needed for adjustment of rare variants (MAF<1%). Therefore, it is possible that admixed or otherwise structured subgroups may be useful in creating a reference similarity map for summary data harmonization provided the reference map contains relevant genetic substructure.

As *Summix2* estimates are dependent on the reference panel, adequate representation is required for accurate substructure estimation and AF harmonization. However, the reference data currently available and used here do not fully represent the genetic diversity within each of the continental groups^65,66^. For instance, the African and South Asian reference populations are only represented by 8 and 5 finer-scale reference groups, respectively, compared to 21 East Asian and 12 European finer-scale reference groups^65-69^. Similarly, the Indigenous American group, with only 47 individuals compared to 704 in AFR, 787 in EAS, 741 in EUR, and 545 in SAS (Supplementary Table 2), is vastly underrepresented by sample size alone^70^. Many consortia, such as the African Diversity Project, All of Us, the GenomeAsia 100K Project, and the Population Architecture through Genomics and Environment (PAGE) study, are pursuing efforts to improve the representation of genetic data from non-European groups^5-8^. While additional representation from diverse ancestries, cultures, and environments is critical to creating generalizable and impactful research, gathering and using data must be done with care. Especially in historically marginalized groups that often see few rewards for genetic studies, such as Indigenous Americans, research must incorporate the values of data sovereignty, community engagement, and trust building^71,72^. Additionally, groups are often labeled (e.g., 1KG, HGDP) according to the geographic region from which the sample originated, which likely does not represent the full genetic diversity of the people. Thus, we follow the National Academies Report recommendation using the “-like” suffix (e.g., 1KG-Pel-like) to indicate similarity to the particular reference group rather than all people^11^.

Until recently, modeling substructure in genetic summary data has been limited, leading to biased results or the underuse of data. This is especially true for genetically admixed populations (often the same groups that are also underrepresented in research)^73^. Here, we show that the methods in *Summix2* can detect, adjust, and even harness substructure in summary data by estimating genetic similarity to reference groups using only summary level data. *Summix2* can detect local and finer-scale genetic substructure as well as substructure beyond ancestry, laying the foundation for better use and harmonization of summary level data. Ultimately, we hope the suite of methods in the *Summix2* software package will enable more robust and equitable use of genetic summary data.

## Online Methods/Methods

### Data

All data was mapped to the GRCh38 build. AFs from gnomAD version 3.1.2^38^ for 644,267,978 variants that passed gnomAD’s quality control (QC) filter for 76,156 whole genomes from ten groups (i.e., AFR/AFRAM, AMI, AMR, ASJ, EAS, FIN, NFE, MID, SAS, and OTH) obtained in November 2021 were used as the observed data. Additionally, we used reference AFs for 76 HGDP & 1KG groups that were harmonized with gnomAD v3.1.2. After removing 24 admixed groups, one group that was not classified in the five continental groupings, and one group with N=1, 50 finer-scale reference groups remained (**Experiment Overview** in **Results;** Supplementary Table 2). We merged the observed gnomAD data and HGDP & 1KG reference data by chromosome, variant position, reference allele, and alternate allele. Variants with missing values in either dataset were removed. We then limited to variants with MAF>0.01in at least one of the 50 finer-scale reference groups or in at least one of the ten gnomAD v3.1.2 groups, resulting in 75,526,117 variants across the 22 autosomes. This data was used for evaluation of finer-scale groups. For comparisons using continental level reference groups, the data was additionally filtered to keep variants within MAF>0.01 in at least one continental reference group resulting in 25,087,586 variants. Continental level AFs were calculated as a weighted sum of the finer-scale reference groups within each continental level group as defined by gnomAD v3.1.2 (Supplementary Table 2).

AFs from the CCPM biobank for 138 of the 148 SNPs from PGS00084^61^ were obtained for individuals classified as European ancestry (for full details on ancestry group assignment see publication^62^). AFs were calculated for individuals with and without prostate cancer, as defined by prostate cancer ICD-9 and ICD-10 codes comprising PheCode 185^74^. ICD codes for cases were 185, 233.4, V10.46, C61, D07.5, Z85.46. Individuals without any of the ICD codes were defined as controls.

Simulation AFs were generated assuming Hardy-Weinberg Equilibrium using the *rmultinom* function in base R with the probability defined using the AFs from the finer-scale or continental reference groups. Simulation code is available on accompanying GitHub site (see **Data Availability**) and more information is in the **Results** under ***Experiment Overview***.

### Summix

*Summix* estimates subgroup mixing proportions (*πk*) by minimizing the LS difference for N AFs in the observed sample (*AF*_*obs,i*_) and AFs from K reference groups (*AF*_*ref,i,k*_) where *i* is the variant indicator, and *k* is the reference group indicator (equation (1)).

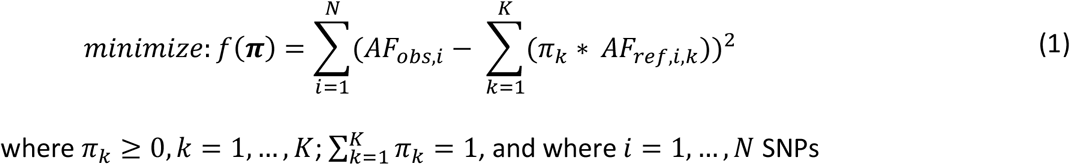

We used sequential quadratic programming (SQP)^75^ to efficiently estimate *πk*. Full details on the original *Summix* method can be found in Arriaga-MacKenzie, et al.^36^

### Generalized Goodness-of-fit Metric

The GoF metric from Summix v1 was simply the LS objective function, *f*(***π***). To identify the relationship between MAF and the GoF metric, we simulated 108,404 AFs from chromosome 22 with 85% AFR-like and 15% EUR-like mixing proportions. We binned the variants into MAF bins (0.0-0.01, 0.01-0.02, 0.02-0.05, 0.05-0.1, 0.1-0.3, 0.3-0.5) and estimated the substructure proportions using variants in each bin separately subsequently obtaining a MAF bin specific GoF value. This was repeated for 1000 replicates (Extended Data Fig. 1).

To correct for the relationship between the GoF metric and MAF bin, we developed a weighted GoF measure, equation (2). To determine the weights, *wl*, we estimated the substructure proportions and obtained the original GoF for seven gnomAD groups by the MAF bins outlined above. For the reference data, we used the five continental groups, rerunning the estimation excluding any reference groups with <2% estimated proportion. We observed similar bin weights across the gnomAD groups, and thus averaged over the gnomAD groups to estimate weights for the new GoF measure, equation (2). We then normalized the GoF in each MAF bin to match the largest GoF value (i.e., in the 0.3-0.5 MAF bin) using bin weights as shown in equation (3).

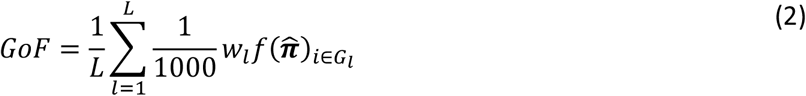

Where there are L MAF bins and 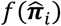 is the LS objective function for SNPs within group, *G*_*l*_. Here, we use L=3 MAF bins of (0, 0.1], (0.1, 0.3], (0.3-0.5].

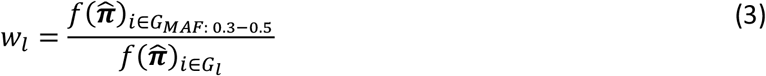

To identify GoF thresholds to indicate poor, moderate, and good fit, we simulated a mixture of five continental reference groups using 1,000 replicates for each simulation scenario. The proportion of one reference group varied between 0.05 to 0.95 by 0.05 increments and was left out of the reference panel. The proportions for the other four groups were randomly assigned to sum to 1 and remained in the reference panel. The process was repeated so that each of the five groups was left out once.

### Substructure adjusted allele frequencies

In the previous release of *Summix*, adjusted AFs were calculated by leaving out a single reference group, *l*, of the user’s choosing. To ensure the adjusted AFs are less dependent on a single reference group, we updated the default method for adjusted AFs to iteratively leave out each reference group, and then average across *K* iterations (equation (4)).

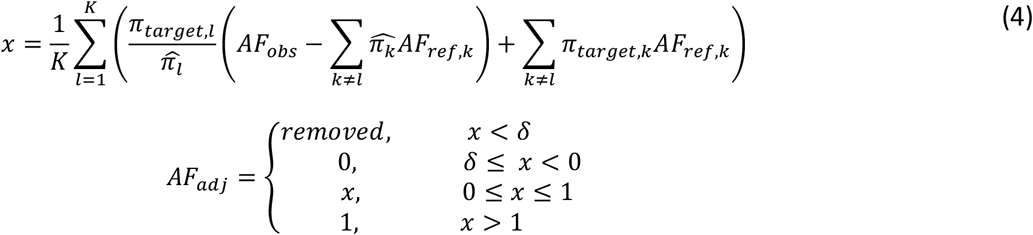

Where *AF*_*adj*_ is the adjusted AF, *l* is the index for the reference group left out, and δ is the precision threshold for which adjusted AF are removed from the dataset or rounded to 0. We found a value of δ = −0.0005 worked well. We tested the performance of the previous AF adjustment method compared to the updated average adjusted AF method across a variety of scenarios (Supplementary Note 6, Supplementary Figs. 7-12). We still enable a leave one out AF adjustment if desired.

Just as the adjusted AFs are a weighted combination of the observed and reference data, the effective sample size for the adjusted AF is also a weighted combination of the observed and reference sample sizes. Thus, *Summix2* now estimates an effective sample size, *N*_*eff,average*_, equation (5), for average adjusted AF using equation (4) and an effective sample size, *N*_*eff,Leave out l*_, equation (6), for adjusted AF calculated using the original *Summix* AF adjustment method, equation (1)^36^. A warning is produced if the effective sample size is less than 50% of the sample size of the observed group.

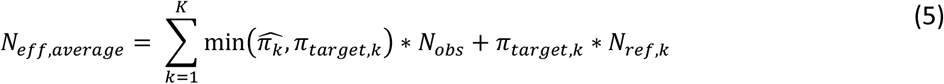

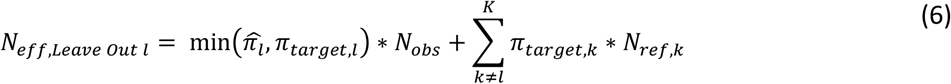

Where, *N*_*obs*_ is the sample size of the observed group and *N*_*ref,k*_ is the sample size of *k*^*th*^ reference group (Details and examples in Supplementary Note 4, Supplementary Table 14).

### Summix-local

We developed and evaluated *Summix-local* for estimating local substructure using two chromosome scanning algorithms: sliding windows, and fast catch-up^43^. For the sliding window algorithm, the user defines the size and overlap of the windows in base pairs (bp) or number of SNPs. *Summix-local* estimates the substructure proportions using the SNPs within each window and averages the substructure proportions within the overlapping sections. By allowing for dynamic window sizes, the fast catch-up algorithm overcomes a limitation in which fixed window size may obscure breakpoints between ancestral segments. The fast catch-up algorithm uses a start-and end-pointer. Beginning at the start-pointer, a window is created using a user-specified number of variants, or size in bp. During each iteration, the end-pointer moves forward one SNP, and the substructure proportions are estimated and compared to the previous iteration (i.e., the window with one fewer SNP included). The algorithm ends and a block is defined when a percent difference in one of the reference groups is larger than the predefined threshold (default 2%). The start-pointer is then advanced to the current end-pointer SNP and the algorithm begins again, continuing until the end of the chromosome is reached. We refer to windows as the input for substructure estimation and blocks as the output. For the sliding window approach, block and windows are the same. For the fast catch-up method, blocks defined as overlapping windows with the same substructure proportions (i.e. substructure proportion difference<2%). Additionally, by default within *summix-local*, we exclude reference groups with <2% global proportions as estimated by Summix (Supplementary Note 7).

We developed a test to identify local chromosomal regions with different substructure proportions compared to the global average, equation (7). We use a Wald test statistic where 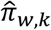 is the substructure proportion estimate for block *w* and reference group *k*, 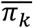 is the average substructure proportion over *W* blocks for the chromosome, excluding block *w*. P-values are estimated using the t-distribution with N – 1 degrees of freedom, where N is the number of SNPs in block *w*. A Bonferroni-corrected threshold of 0.05 correcting for the total number of windows (*W*) and K-1 reference groups is used to adjust for multiple testing.

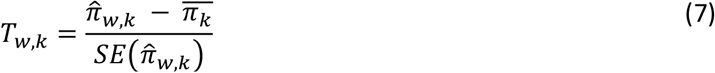

To estimate the standard error, 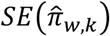, we average estimates of within block and between block error for each reference group, *k*. We use simulations to estimate within block standard error.

Briefly, we create 1000 replicates of simulated reference AFs using a multinomial distribution with parameters equal to the number of haplotypes and AFs in the real reference data. The simulated reference AFs are used to estimate the within block substructure proportions of the observed data. The within block standard error estimate is the standard deviation of the resulting sampling distribution. To account for between block variability, we estimate the standard deviation of the proportion estimates over the chromosome.

### Local substructure simulations

We filtered the gnomAD v3.1.2 chromosome 19 data for LD using PLINK2.0, removing variants with a pairwise r^2^>0.2 using *–indep-pairwise 300 100 0*.*2*. We simulated SNP genotypes for both observed and reference samples using *rmultinom*, assuming Hardy-Weinberg equilibrium, and using reference AFs as the probabilities. We simulated AFR/AFRAM-like observed data with 85% AFR-like and 15% EUR-like proportions, and AMR-like observed data with 10% AFR-like, 65% EUR-like, and 25% IAM-like proportions. We assessed observed sample sizes of 1000, 10,000*, and 50,000 individuals, reference samples of 100, 500*, and 1000 individuals per group, and window sizes of 200, 1,500*, and 3,000 variants. The default simulation parameters are marked by an asterisk. We changed one parameter type keeping the other parameters at default values to evaluate a variety of simulation scenarios. We assessed minimum accuracy and precision of the substructure estimates across the blocks using equations (8) and (9).

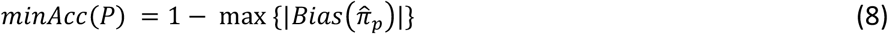

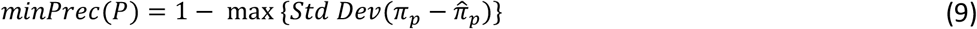

Where 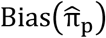 is the mean difference between the simulated proportion, π_p_, and the estimated proportion of genetic substructure, 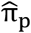, and P is the number of tests, here the number of blocks (W).

To assess type I error and power to detect regions where the local substructure differs from the average chromosomal global substructure, we replaced a segment of the simulated observed data with different mixing proportions using a magnitude of change of 2%, 3%, or 5% and region size of 50Kb, 400Kb, or 1,000Kb. The default parameter scenario had a region size of 400kb with a 3% difference in IAM-like mixing proportion for the AMR-like simulation scenario and 2% difference in AFR-like mixing proportion in the AFR/AFRAM simulation scenario. We defined type I error as the proportion of simulations out of the 100 replicates with at least one significant block outside of the simulated signal region (i.e., in the null regions of the chromosome). Similarly, we defined power as the proportion of simulation replicates with at least one significant block overlapping the simulated signal region.

### Local-substructure scan in gnomAD

We applied *Summix-local* to gnomAD AMR and AFR/AFRAM using the fast-catchup algorithm and the default parameters. For computational efficiency, we assessed each chromosome separately, using the merged dataset that was filtered for continental reference MAF and the continental reference AF data. We removed blocks with poor fit (GoF > 1.5) from further analysis. Multiple testing correction was accounted for by a Bonferroni-correction for 5945 genome-wide blocks and the number of reference groups (two and three for AFR/AFRAM and AMR, respectively) resulting in a threshold of 4.21e-6 and 2.80e-6.

To identify previously putative signals of selection, we performed a literature search on April 30^th^, 2023, using Google Scholar for studies of signatures of selection in Latino and African American populations by searching “selection in humans” (Supplementary Table 5). We then selected the blocks that overlapped by bp position with these previously identified regions. We define replication as the overlap of a block by at least one SNP with a significant p-value after multiple testing correction for the number of reference groups and the number of replication signals being tested. This resulted in significance thresholds of 1.56e-3 for AFR/AFRAM (2 reference groups and 16 regions) and 5.75e-4 for AMR.

We performed gene set enrichment for the genome-wide significant blocks using the Gene Ontology (GO) Enrichment Analysis^58,59^ tool (https://www.geneontology.org/) searching the PANTHER GO-Slim Biological Process^76^. PANTHER-GO is a comprehensive resource for gene classification according to functional classifications using the GO. The ‘slim’ annotation set includes only a subset of the GO, which have been curated to be the most informative GO terms to infer gain or loss of function and are evolutionarily conserved. We selected this subset to identify the most likely pathways under selection and to ease interpretation. We identified genes with overlap or fully contained in the significant blocks using the UCSC Genome Browser Table Browser tool^77^. The GO Enrichment Analysis compares the input gene list to the reference list of all genes in the PANTHER database using Fishers exact test, for each Gene Ontology. The Benjamini-Hochberg False Discovery Rate is used to adjust for multiple testing.

### Fixation index

To estimate the similarity between subgroups, we used the Hudson pairwise Fixation Index (*F*_*ST*_)^42^ to estimate similarity between reference groups, equation (10).

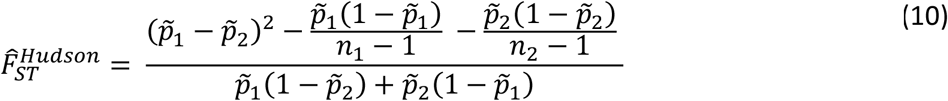

Where *ni* is the sample size and 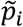 is the sample AF for group *i* for *i* ∈ {1, 2}.

The Hudson pairwise *F*_*ST*_ estimator measures population similarity by averaging the Weir and Hill *F*_*ST*_78 estimates for two groups. The Weir and Hill *F*_*ST*_ estimator calculates the correlation between alleles from a given population relative to its most recent ancestral population. Unlike other *F*_*ST*_ estimators, estimates made by 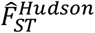 are not sensitive to the ratio of samples sizes in each population^42^.

We estimated the *F*_*ST*_ between each pair of the 50 finer-scale reference groups from 1KG and HGDP using individual level data. For estimation, we removed individuals labeled as related in gnomAD v3.1.2 metadata^38^ and used variants with MAF ≥ 0.01 and pruned variants so that pairwise *r*^2^ < 0.2 using *bcftools +prune*. We calculated pairwise *F*_*ST*_ using the 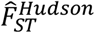 function in *PLINK 2*.*0*^*79*^.

### Finer-scale substructure simulations

We performed simulations using the 50 finer-scale groups as reference data to assess the accuracy and precision of *Summix2* to estimate finer-scale substructure. AFs were simulated independently for each SNP using a multinomial distribution (*rmultinom* in base R), assuming Hardy-Weinberg equilibrium, and using reference AFs as the probabilities. We simulated reference group sample sizes of 100 for each group and observed group sample sizes of 10,000. We used 100 simulation replicates, sampling 100,000 genome-wide SNPs in each replicate. Minimum accuracy and precision are defined in equation (8) and equation (9), respectively, where p is the number of finer-scale reference groups (most often 50).

To evaluate how similarity in substructure affected accuracy and precision of the proportion estimates, we simulated observed data from reference groups with varying levels of pairwise *F*_*ST*_. We simulated three mixture proportion scenarios (i.e., four reference groups simulated at equal, varying, and equal with one zero mixing proportions) across three pairwise *F*_*ST*_ scenarios (pairwise *F*_*ST*_= 0.009, 0.007, 0.005 for high, medium, and low similarity respectively) using EAS finer-scale reference groups. Two-pairs of finer-scale ancestries with the given pairwise *F*_*ST*_ were used as reference ancestries in each simulation scenario. Oroqen and Daur (pairwise *F*_*ST*_=.009) and Yizu and Naxi (pairwise *F*_*ST*_=.009) were selected for the high similarity group, Tujia and Han (pairwise *F*_*ST*_=.007) and Xibo and Mongola (pairwise *F*_*ST*_=.007) were selected for the medium similarity group, and Han Chinese (Chb) and Kinh (Khv) (pairwise *F*_*ST*_=.005) and Tu and Mongola (pairwise *F*_*ST*_=.005) were selected for the low similarity group. See Supplementary Table 9 for simulated reference group proportions.

Using the 50 finer-scale reference groups, we evaluated substructure estimation in real data for gnomAD v3.1.2’s ten groupings: African/African American, Amish, Latino/Admixed American, Ashkenazi Jewish, East Asian, Finnish, Non-Finnish European, Middle Eastern, South Asian, Other. Ninety-percent CIs for the proportion estimates were calculated by randomly sampling 100,000 genome-wide SNPs 100 times (Supplementary Table 10).

### Adjusted allele frequencies

On chromosome 19, we assessed adjusting the observed gnomAD AMR AFs to three target samples: 1) 1KG Pel, 2) a simulated target with continental substructure (10% AFR-like, 15% EUR-like, 75% IAM-like), and 3) a simulated target with finer-scale substructure (45% HGDP-Maya-like, 10% HGDP-Karitiana-like, 10% HGDP-Pima-like, 10% HGDP-Colombian-like, 5% 1KG-Esn-like, 5% 1KG-Yri-like, 8% 1KG-Gbr-like, 7% 1KG-Ibs-like). The proportions used to simulate the continental and finer-scale target samples match those estimated from the 1KG-Pel-like sample using *Summix2*. We estimated the observed reference group proportions using *Summix2* with either continental or finer-scale reference data and at either the global or local level resulting in four estimation groups: 1) continental global, 2) continental local, 3) finer-scale global, 4) finer-scale local. We compared these AF adjustments to adjustments using the local substructure proportion estimates provided by gnomAD v3.1 for the AMR group, which were estimated using continental reference groups (AFR, EUR, and IAM) and RFMix on individual level data^80^. We limited our comparison to SNPs with MAF>0.01 in the target sample with gnomAD local substructure estimates available (i.e., 12,513 out of 234,633 SNPs with MAF>0.01 on chr 19). We calculated the absolute and relative difference between the observed (adjusted or unadjusted) AF to the target AF across MAF bins ([0.01, 0.02), [0.02, 0.05), [0.05, 0.10), [0.10, 0.30), [0.30, 0.50]) and assessed whether the differences varied by adjustment group using paired t-tests. We assessed concordance between the AF for the target sample and the observed sample adjusted AFs using Lin’s CCC.

### Estimating genetic similarity to case-status

We investigated the ability of *Summix2* to detect genetic substructure other than genetic ancestry by evaluating the ability to estimate the proportion of individuals with a genetic similarity to case status (i.e., the genetic predisposition to disease). We used AFs from prostate cancer cases and controls from a 2018 prostate cancer GWAS used to create a prostate cancer PGS (PGS000084) as the reference data to estimate the proportion of people in the CCPM Biobank v2 data that were genetically similar to the prostate cancer cases^61,62^. We derived the case AFs from the reported odds ratios and control AFs (Supplementary Note 5). Out of the 147 SNPs included in the prostate cancer PGS, 138 were in the CCPM Biobank and were used for further analysis. To evaluate estimation across a variety of characteristics, we subset the CCPM Biobank data by biological sex (i.e., female, male), males with prostate cancer, and age group (i.e., < 40 years, 40-60 years, >60 years). We estimated jackknife 95% CIs.

## Data availability

The gnomAD v3.1.2, 1KG and HGDP data is available for download at https://gnomad.broadinstitute.org/downloads. Code used to merge gnomAD v3.1.2, 1KG, and HGDP data and generate simulated datasets as well as an exemplar simulated AF dataset is available on GitHub at https://github.com/hendriau/Summix2_manuscript. The exemplar simulated dataset contains 234,633 LD 0.2 filtered variants from chromosome 19 with observed finer-scale simulated substructure proportions of 45% HGDP-Maya-like, 10% HGDP-Karitiana-like, 10% HGDP-Pima-like, 10% HGDP-Colombian-like, 5% 1KG-Esn-like, 5% 1KG-Yri-like, 8% 1KG-Gbr-like, 7% 1KG-Ibs-like. Fifty simulated finer-scale references and 5 continental simulated references for n = 200 and 500 respectively are included. This simulation is based on the dataset filtered for MAF>0.01 in at least one continental reference group.

## Code availability

*Summix2* software (R package) is available at https://bioconductor.org/packages/release/bioc/html/Summix.html. R software is available at https://www.r-project.org/. PLINK 1.9 software is available at www.cog-genomics.org/plink/1.9/.

## Acknowledgements

We thank the participants of the Colorado Center for Personalized Medicine Biobank. None of this research would be possible without publicly available data from gnomAD, the 1000 Genomes Project, and the Human Genome Diversity Project. This work was supported by the National Human Genome Research Institute (R35HG011293, R01HG011345, U01HG011715). This work also used the computing resources at the Center for Computational Mathematics, University of Colorado Denver, including the Alderaan cluster, supported by the National Science Foundation award OAC-2019089.

## Ethics declarations

Here, we have used publicly available resources from gnomAD and use the gnomAD v3.1.2, 1KG, and HGDP group labels^38^. We recognize that group labels are somewhat arbitrary and are often based on geographic location of the individuals included. For consistency, we use the assigned labels when directly referencing the public data. When we report estimated or simulated genetic substructure similarity proportions, we have adopted the “-like” nomenclature as recommended by the National Academies 2023 report^11^. Additionally, we use the group label Latino for people whose origin are in Latin America rather than Latinx, as Latinx is a gender-neutral term primarily used by English-speaking people in the U.S. with a research-based designation, rather than community based term^81^.

We use CCPM Biobank data for individuals classified as European ancestry using genotype information (for full details see publication^62^). We recognize that restricting research to individuals of European ancestry is not optimal and limits generalizability of the research conclusions. In future research, we will evaluate estimating and harmonizing over multiple levels of genetic substructure (e.g., genetic admixture and study ascertainment).

