## Extended Data Figs. 1-8 for "Characterizing substructure via mixture modeling in large-scale genetic summary statistics"

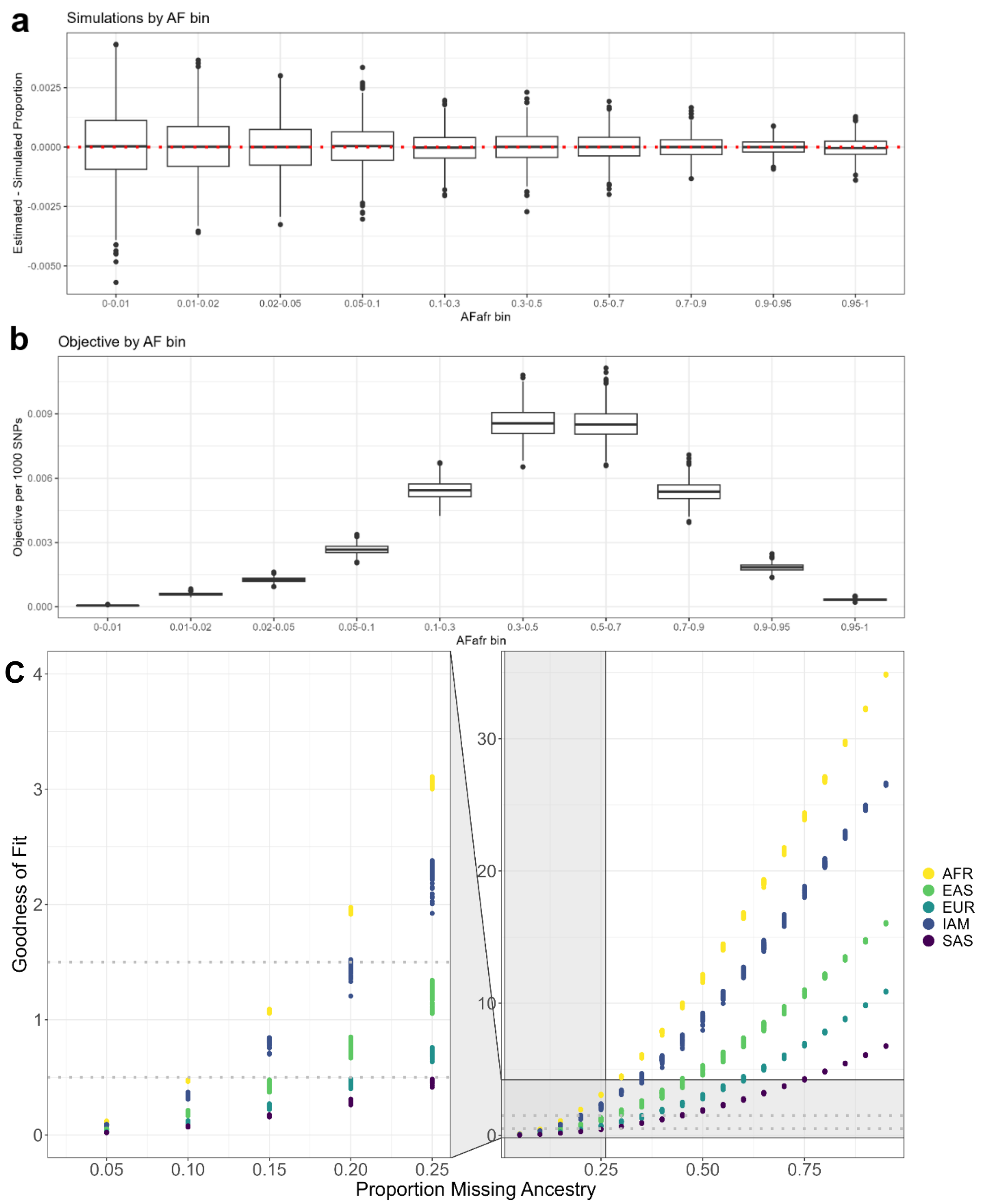


**Extended Data Fig. 1: Simulations show original Summix goodness of fit measure varies by allele frequency**. We used simulations to explore the relationship between by AF bin (x-axis) and the estimated substructure proportion accuracy (y-axis) **a,** and goodness of fit (GoF; y-axis) **b**. While accuracy is consistent across the AF bins, the original GoF measure decreases with decreasing MAF bin resulting in a lack of generalizability of the GoF measure to other genotyping technologies with different AF distributions. We developed a weighted goodness of fit measure adjusting by AF bin. **C,** Simulations were used to determine good, moderate, and poor fit thresholds for GoF. We simulated one reference group at a fixed proportion, and randomly varied the remaining four groups to sum to 1. We then estimated the substructure proportions using *Summix2* without the fixed group in the reference panel. We used 1000 replicates per simulation scenario, varying the reference group left out and the proportion missing. Resulting GoF thresholds where <0.5 for good fit, 0.5 - 1.5 for moderate fit, and >1.5 for poor fit (dotted horizontal lines).

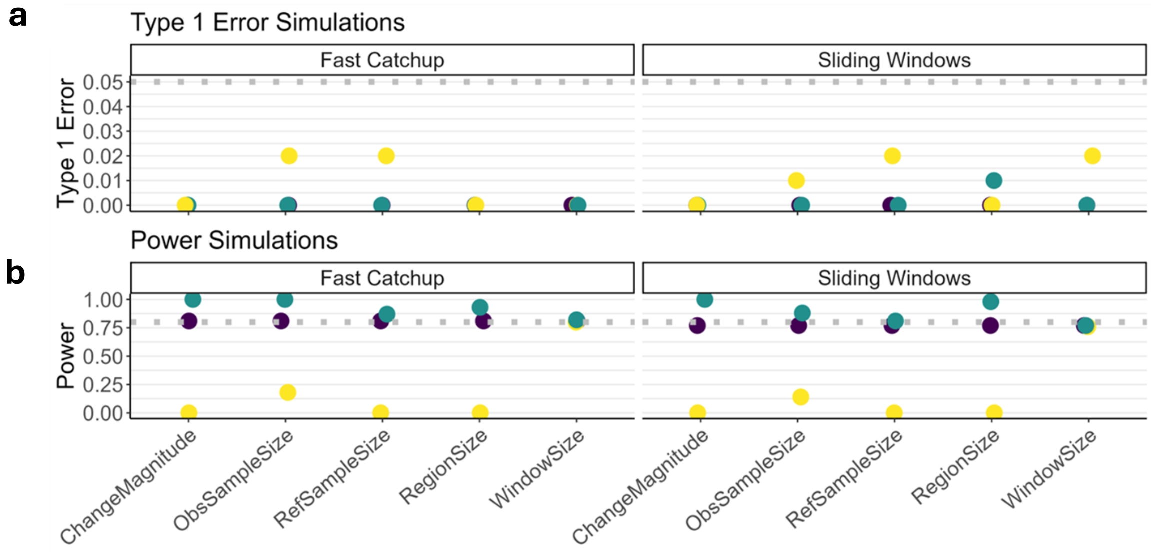


**c**

| Parameter | Small | Default | Large |
|---|---|---|---|
| Change Magnitude | 1% | 2% | 5% |
| Observed Sample Size | 1,000 individuals | 10,000 individuals | 50,000 individuals |
| Reference Sample Size | 100 | 500 | 1000 |
| Region Size | 50,000 bp | 400,000 bp | 1,000,000 bp |
| Window Size | 700 variants | 1500 variants | 2500 variants |

**Extended Data Fig. 2**: **Summix-local detects regions where local substructure differs from global in an AFR/AFRAM-like sample a,** Type I error and **b,** power for local substructure scan for fast-catchup (left) and sliding windows (right) algorithms across simulation scenarios as shown in **c**, for an AFR/AFRAM-like simulated observed sample. LD filtered chromosome 19 data for 234,633 variants was used to simulated 80% AFR-like and 20% EUR-like substructure, with a region (default 400,000bp) was replaced with simulated data with differing substructure proportions (default 82% AFR-like, 18% EUR-like). Type I error is maintained below a 5% threshold for all scenarios while power is at or above an 80% threshold for all scenarios other than the small parameters (yellow). Power and type I error are similar between the fast catchup and sliding windows algorithms.

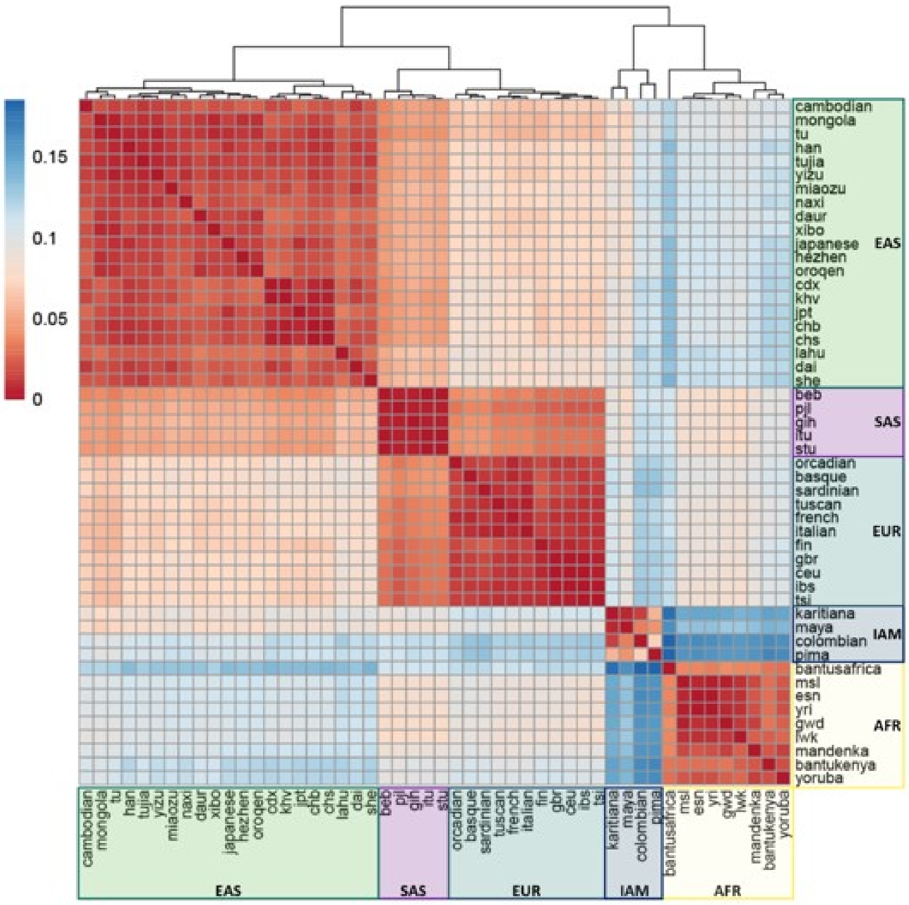


**Extended Data Fig. 3: Comparison of genetic similarity within continental and finer-scale reference groups.** Pairwise $F_{ST}$ ranging from .001 (red, indicating high similarity) to .035 (blue, indicating low similarity). Fifty finer-scale groups within continental reference groupings as defined by 1KG and HGDP: EAS (East Asian, green), SAS (South Asian, purple), EUR (European, teal), IAM (Indigenous American, blue), AFR (African, yellow).

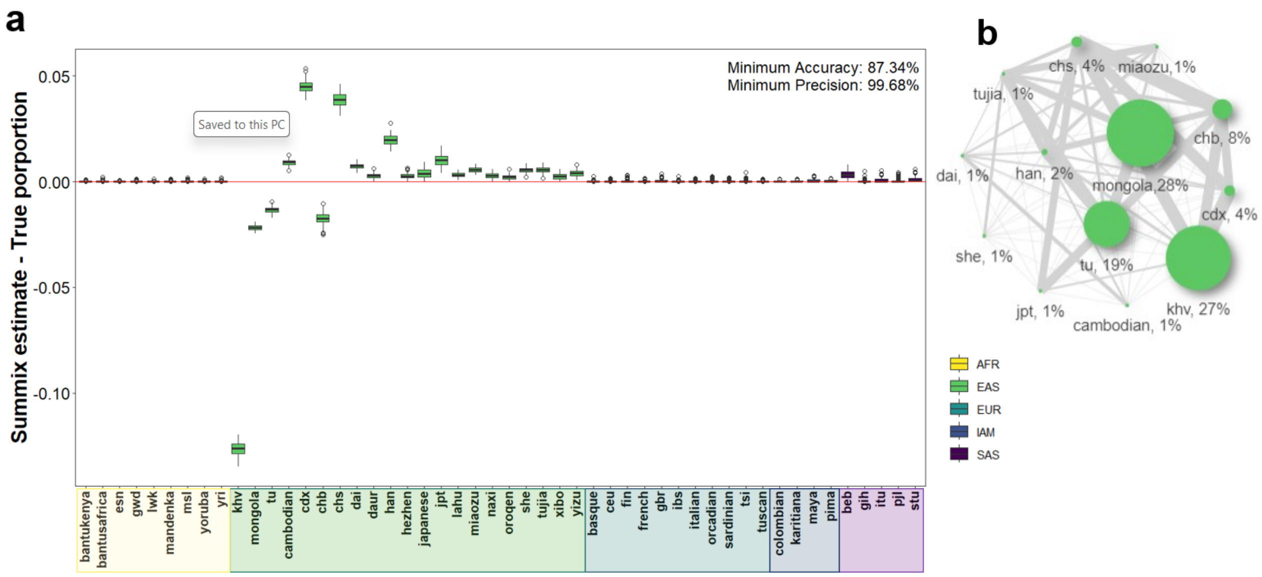


**Extended Data Fig. 4: Bias from *Summix2* estimates is contained in larger subgroup and can be captured by genetic similarity mapping. a**, Accuracy (y-axis) and precision for EAS-like population ($\pi_{chb}$=0.1, $\pi_{tu}$=0.2, $\pi_{mongola}$=0.3, $\pi_{khv}$=0.4, pairwise $F_{ST}$ =.005 grouping, 'Varying mixing proportions', Supplementary Table 9). Maximum bias is 0.127 but is contained within the larger East Asian continental reference grouping. **b**, Genetic substructure similarity map of *Summix2* estimates for EAS-like population where edge thickness between nodes indicates pairwise similarity (thicker edges indicate higher similarity) as defined by pairwise $F_{ST}$ and node size indicates magnitude of the *Summix2* mixture proportion estimate for the given reference group.

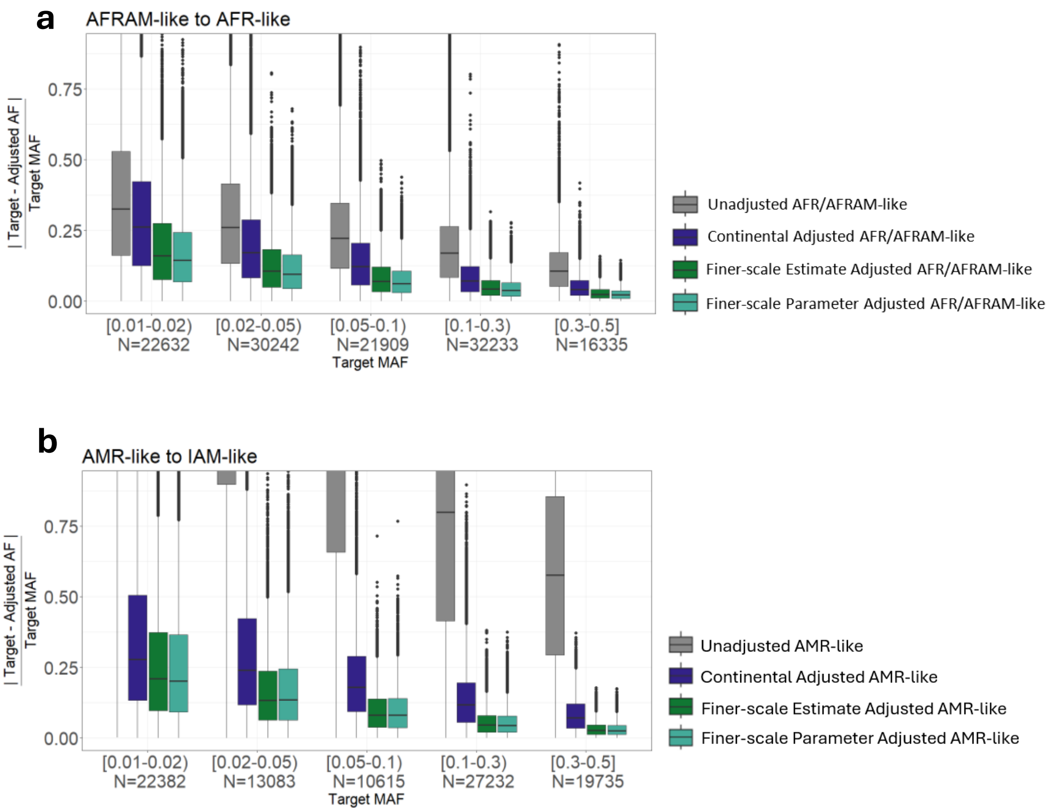


**Extended Data Fig. 5. Summix adjusts for finer-scale genetic substructure.** Relative difference between target AFR-like sample AFs and unadjusted (gray) or adjusted observed AFR/AFRAM-like AFs grouped by MAF bin (continental adjustment, blue; finer-scale estimate adjustment, green; finer-scale parameter adjustment, teal). **b**, Relative difference between target IAM-like sample AFs and unadjusted (gray) or adjusted observed AMR-like AFs grouped by MAF bin (continental adjustment, blue; finer-scale estimate adjustment, green; finer-scale parameter adjustment, teal). Simulated proportions for (**a**) and (**b**) in Supplementary Table 11.

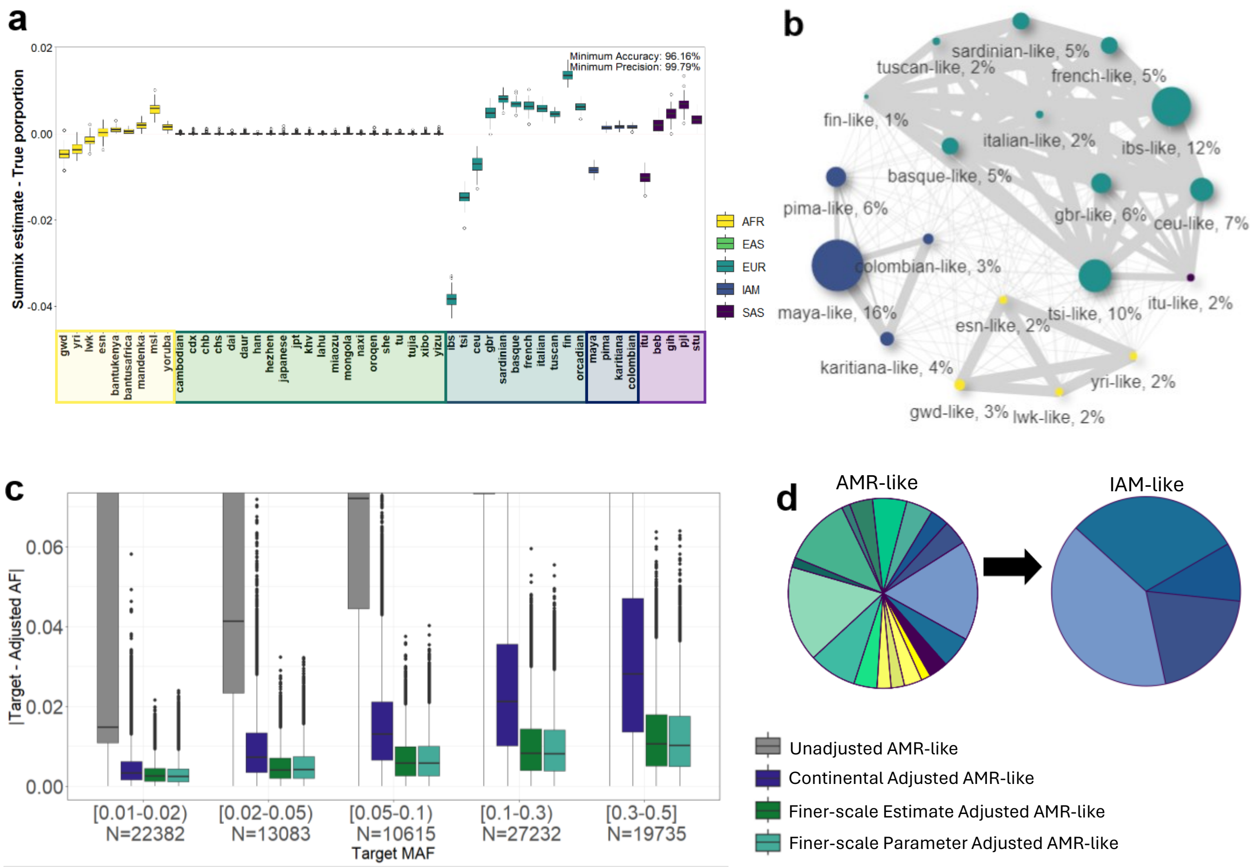


**Extended Data Fig. 6. Summix2 detects and adjusts for finer-scale genetic substructure.** **a**, Accuracy of AMR-like simulation scenario. **b**, Genetic substructure similarity map of Summix2 estimates for the AMR-like sample where edge thickness between nodes indicates pairwise similarity (thicker edges indicate higher similarity) as defined by pairwise $F_{ST}$ and node size indicates magnitude of the *Summix2* mixture proportion estimate for the given reference group. **c,** Absolute difference between target AMR-like sample AFs and unadjusted (gray) or adjusted observed IAM-like AFs grouped by MAF bin (continental adjustment, blue; finer-scale estimate adjustment, green; finer-scale parameter adjustment, teal). **d**, Finer-scale substructure proportions for AMR-like sample and IAM-like sample in (d); simulated proportions in Supplementary Table 11. (Note: “-like” nomenclature omitted from x-axis plot labels for simplicity).

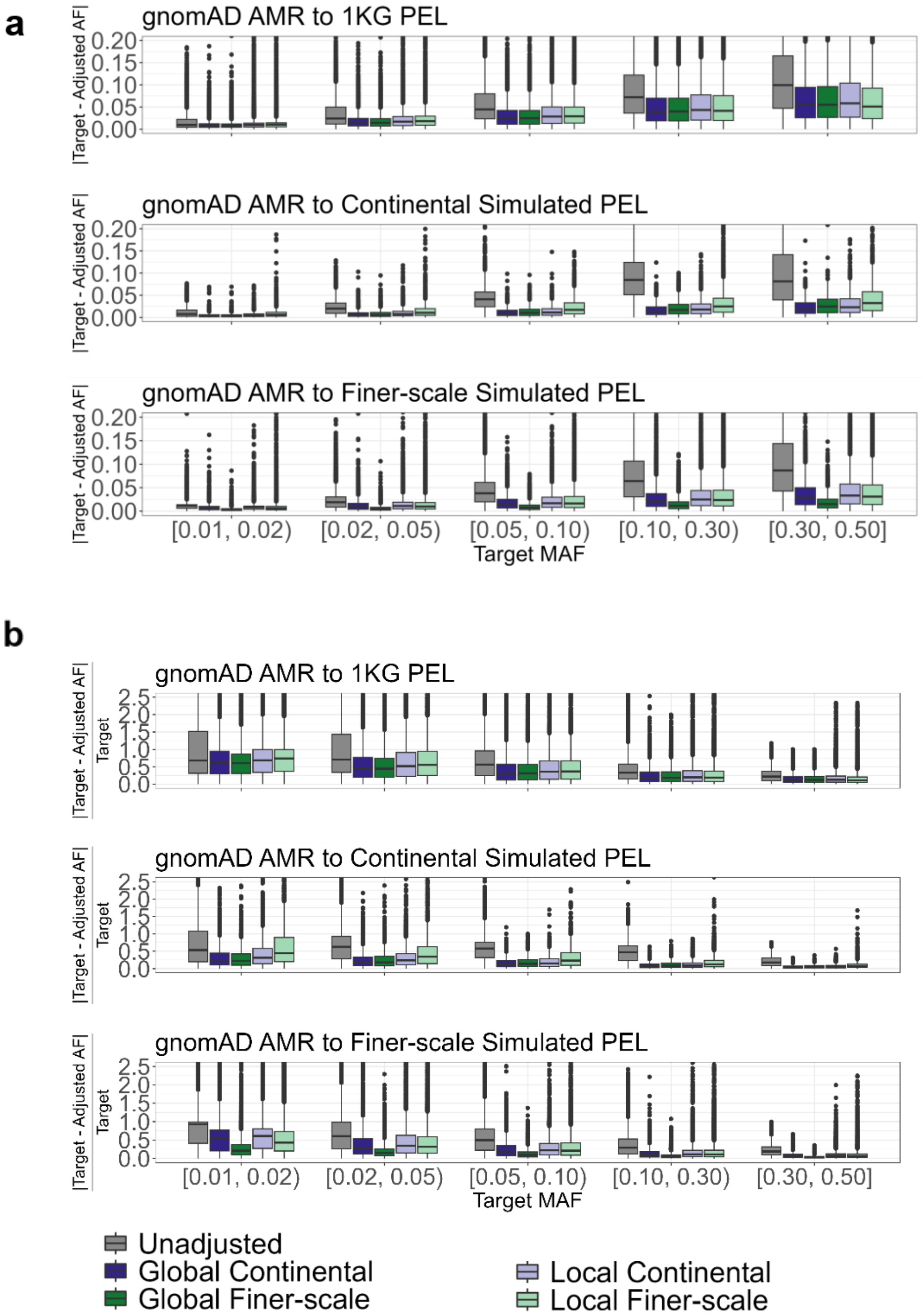


**Extended Data Fig. 7: *Summix2* estimates enable harmonization for >15x variants**. The absolute (**a**) and relative (**b**) difference for AF harmonization of gnomAD AMR to three different target samples for **top,** 1000 Genomes Peruvian **middle,** simulated Peruvian-like with continental-level substructure **bottom,** simulated Peruvian-like with fine-scale substructure, by minor allele frequency bin (x-axis) using all 234,633 LD pruned variants on chromosome 19 (compared to 12,513 variants with local substructure information from gnomAD). Further details are in the Results, and **Online Methods**.

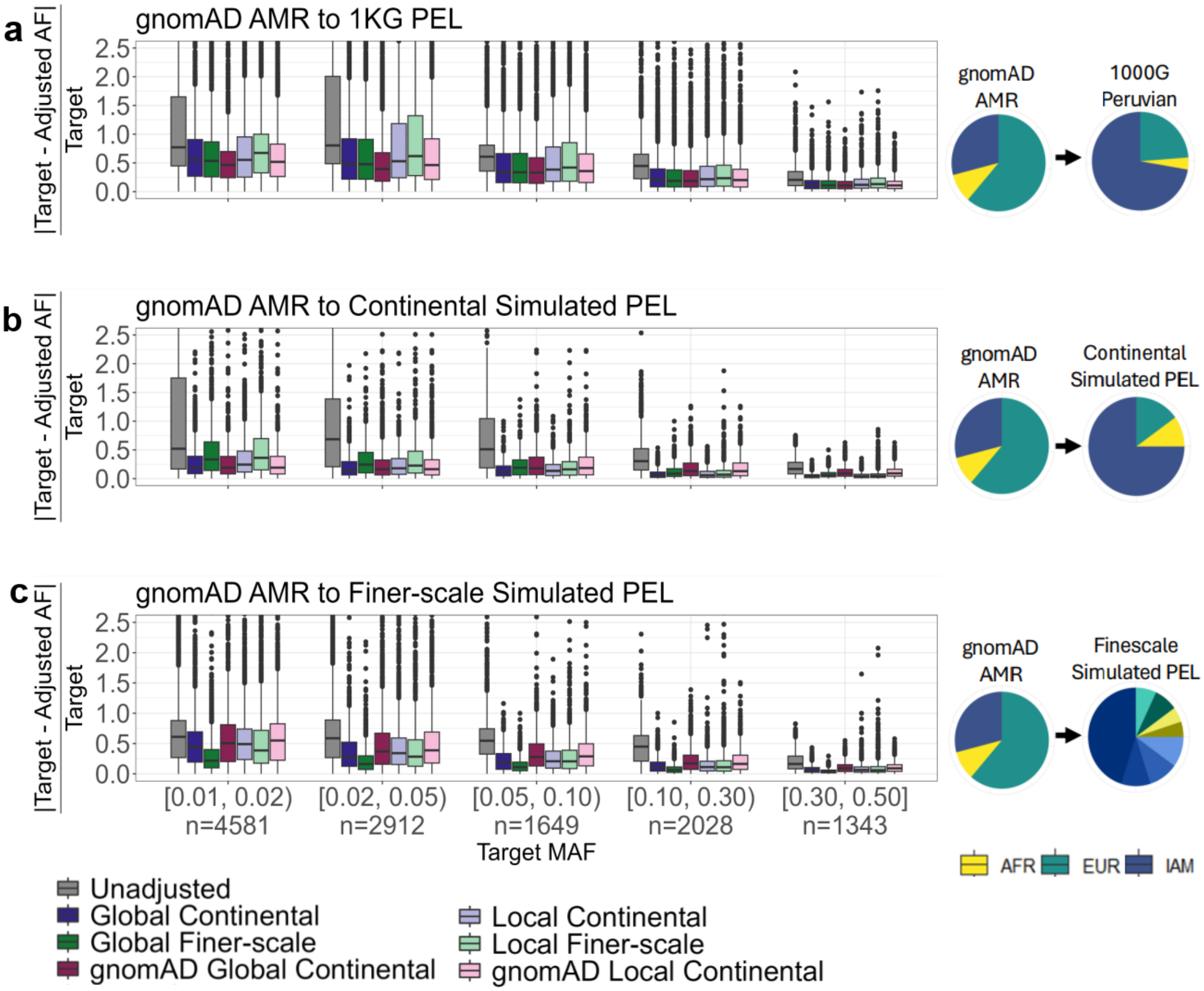


**Extended Data Fig. 8: Allele frequency adjustment improves data harmonization.** Relative difference between the target AF and observed AF (y-axis) with or without adjustment of the gnomAD AMR sample to match three different target samples **a,** 1000 Genomes Peruvian **b,** simulated Peruvian-like with continental-level substructure **c,** simulated Peruvian-like with fine-scale substructure, by minor allele frequency bin (x-axis). Results for 12,513 variants on chromosome 19 with gnomAD local substructure estimates, and filtering for LD ($r^2 > 0.2$) and MAF (>1%). Unadjusted gnomAD AMR (grey) is always outperformed by the adjusted AFs using gnomAD (global – red, local – pink) and Summix2 (global continental – blue, global finer-scale – green, local continental – light blue, local finer-scale – light green). Global substructure proportions for observed and target data is in pie charts (right).
