## Supplementary Information for "Characterizing substructure via mixture modeling in large-scale genetic summary statistics"

### Supplement

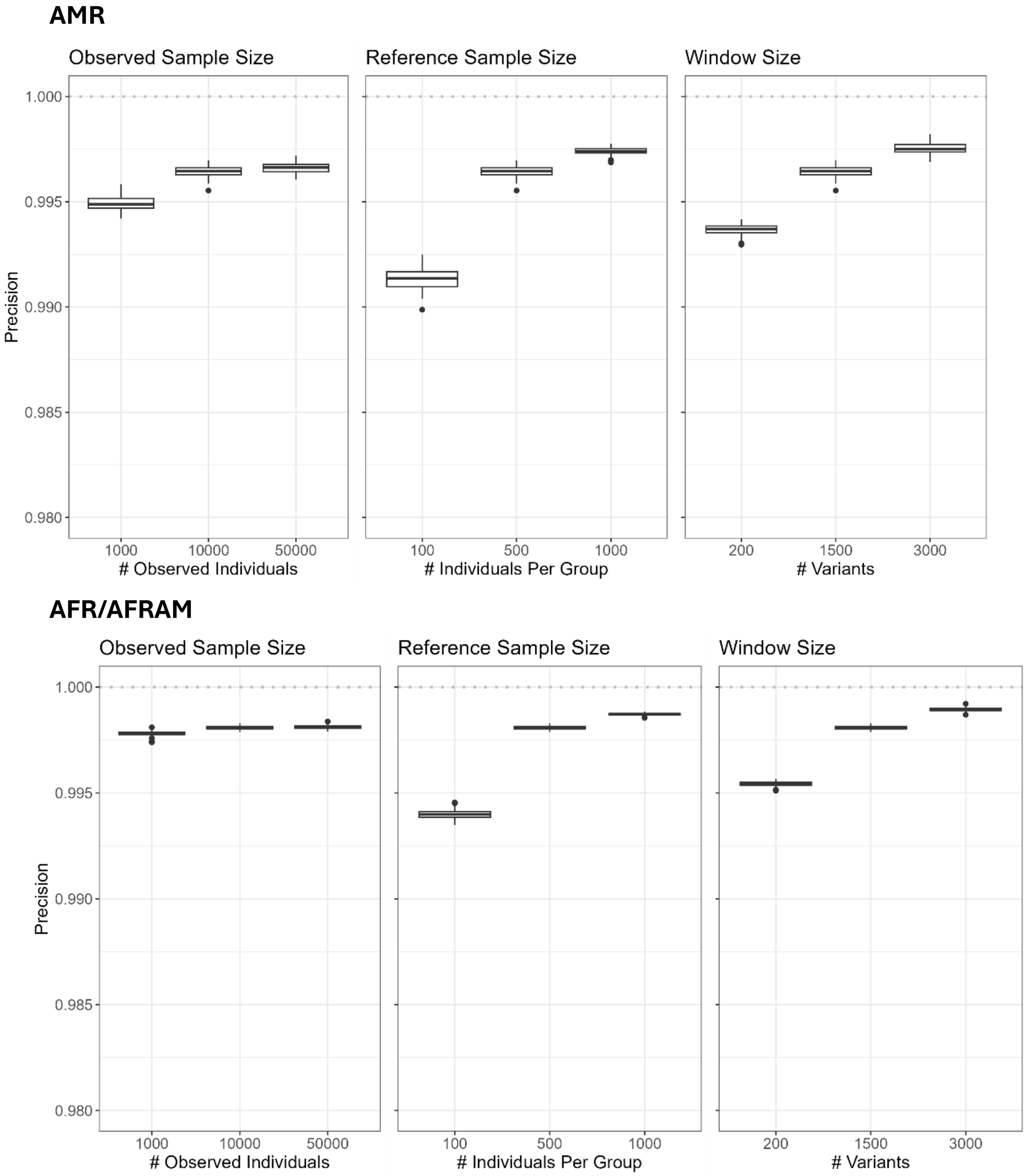


**Supplementary Fig. 1**: **While high for all scenarios, minimum precision for *summix-local* estimates decreases with sample size and number of variants.** Minimum precision for chromosome 19 local substructure blocks for 100 simulation replicates varying observed sample size (left column), reference group sample size (middle column), and window size (right column). Unless changing (i.e. x-axis), default parameters (N observed = 10000, N reference = 500 per group, window size = 1500 variants) are used. The AMR-like sample (top) was simulated to have 65% EUR-like, 25% IAM-like, and 10% AFR-like proportions, while AFR/AFRAM-like (bottom) were 80% AFR-like and 20% EUR-like.

**Supplementary Note 1. Correlated variants affect local substructure estimate precision, but not accuracy.**

In order to assess the effect of LD on the accuracy and precision of the *Summix-local* substructure estimates, simulations were used to model correlation between SNPs. We pruned the chromosome 19 data for LD using PLINK 1.9[74], removing any SNPs with pairwise $r^2 > 0.2$ or 0.8. We then simulated a sample with 85% AFR-like and 15% EUR-like proportions. To simulate perfect correlation, each of the 100 SNPs in a window was doubled once, twice, or three times, and the sliding windows algorithm was applied on those windows of 100, 200, 400 or 800 variants, respectively, so that each window had 100 effective number of SNPs. The window overlap was 20 variants. We find that perfect correlation of SNPs within a window does not affect the accuracy of the substructure proportion estimates. Additionally, we compared the accuracy and precision for the *Summix-local* sliding windows algorithm on windows of 100 SNPs with different LD pruning thresholds ($r^2$ 1.0, 0.8, 0.2). Each region contained 100 SNPs post filtering. The LD pruned data resulted in higher variability (i.e. lower precision) in local substructure estimates, with minimum precision decreasing from 97.5% to 95%; however accuracy remained high (>98%). We used LD pruning (removing SNPs with $r^2$>0.2) in subsequent analyses to more accurately represent the number of independent SNPs in each window (Supplementary Fig. 2).

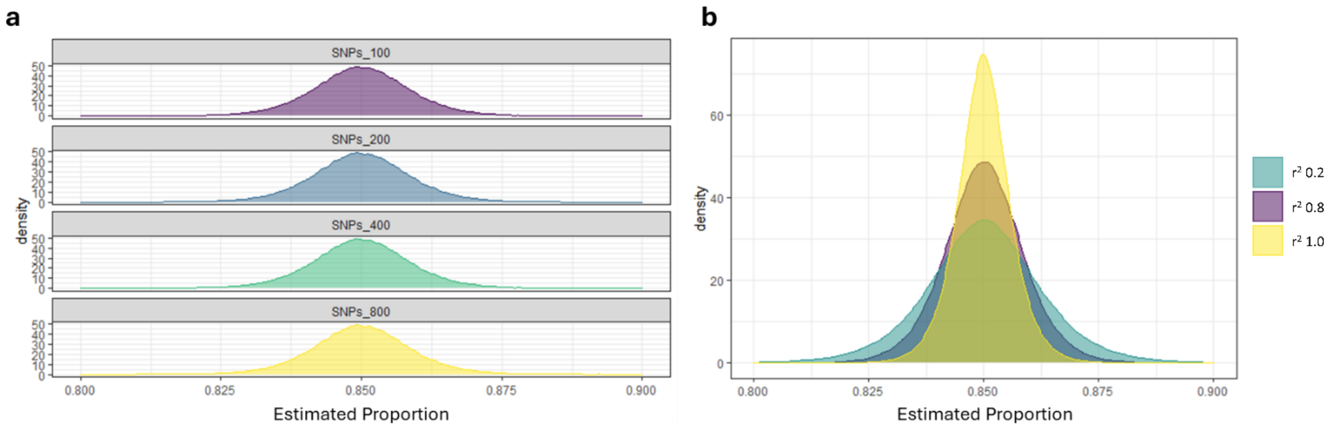


**Supplementary Fig. 2: Accuracy of *Summix* estimates is stable by LD structure, while the effective number of independent SNPs decreases with higher LD.** AFR-like proportion *Summix* estimates simulated at a value of 0.85 with 1000 replicates **a,** Distribution of estimates for 100 independent SNPs either not duplicated (N=100, purple), duplicated once (N=200, blue), duplicated twice (N=400, green), or duplicated four times (N=800, yellow). **b,** Distribution of substructure proportion estimates using LD pruned and non-pruned datasets ($r^2$ 1.0 yellow, 0.8 purple, and 0.2 green) with the same total number of variants per window (N=100). While the accuracy is constant, the variability is smallest when not LD pruned ($r^2$ =1.0) and greatest for stricter LD pruning ($r^2$ =0.2), since the estimates are using correlated variants, thus reducing the variability.

**Supplementary Note 2. Precision of local substructure block boundaries.**
We assessed the precision of *summix-local* for detecting the end points of the simulated signal shown in Extended Data Fig. 2, where gnomAD chr19 v3.1.2 LD filtered data (234,633 SNPs) was used to simulate a chromosome with 85% AFR-like and 15% EUR-like substructure proportions. Then a region of varying sizes was simulated to instead have 87% AFR-like and 13% EUR-like substructure proportions. Over 100 simulation replicates, we determined the start and end of the blocks by relative position in the chromosome (first SNP is 1, the last SNP is 234633) and compared it to the true start and end position of the region with different substructure proportions. We used different window size parameters and compared the fast catch-up and sliding windows algorithms. Note, often multiple consecutive blocks were significant. When this was the case, we merged them into a single significant block for this analysis (in these cases the mean number of SNPs in the significant block will be larger than the window size parameter).

In the scenarios evaluated, the detected block begins before the true region begins. In many cases, the blocks also ends before the true region ends (Tables 1 and 2). Exceptions for this are generally when the window size is larger than the true region size. On average, the fast catch-up algorithm had smaller mean distance from the true block coordinates than the sliding windows algorithm (598 vs 634 SNPs from the start position; 84 vs 326 SNPs from the end position). Exceptions include large mean number of SNPs from the start coordinate when the region size was 400,000bp and the window size was 2500-3000 variants. This is because the minimum window size was much larger than the true region to detect. For the sliding windows algorithm, the maximum number of SNPs from the true start of the block occur is the window size (Table 3). Thus, assuming a region has been detected, the maximum distance from a start or end for the sliding windows algorithm is the window size minus one base pair for the overlap. The mean distance is half of the window size. Overall, the fast catch-up algorithm achieves more or similar precision in the detected region coordinates compared to the sliding windows algorithm. Additionally, better precision is achieved when the window size is smaller, but at the cost of computational efficiency, as runtime is linear with the total number of windows.

**Table 1. Difference in coordinates for significant blocks using fast catch-up algorithm**

| Region Size | | Window Size (SNPs) | | Mean* # SNPs from | | Mean # SNPs in block |
|---|---|---|---|---|---|---|
| bp | SNPs | Min | Max | start | end | |
| 400,000bp | 1086 | 1500 | 2000 | -573 | -145 | 1513 |
| 400,000bp | 1086 | 700 | 1000 | -54 | -133 | 1006** |
| 400,000bp | 1086 | 2500 | 3000 | -1178 | +241 | 2504 |
| 1,000,000bp | 2729 | 1500 | 2000 | -588 | -298 | 3018** |
| | | | Average | -598 | -84 | |

*Mean is over 100 replicates
**When output contained multiple consecutive significant blocks, they were merged for this analysis, resulting in blocks with total number SNPs in block greater than max window size

**Table 2. Difference in coordinates for significant blocks using sliding windows**

| Region Size | | Window | | Mean* # SNPs from | | Mean # SNPs in block |
|---|---|---|---|---|---|---|
| bp | SNPs | Size | Overlap | start | end | |
| 400,000bp | 1086 | 2000 | 200 | -925 | -84 | 2000 |
| 400,000bp | 1086 | 1000 | 200 | -38 | -26 | 1000 |
| 400,000bp | 1086 | 3000 | 200 | -577 | +1337 | 3000 |
| 1,000,000bp | 2729 | 2000 | 200 | -994 | +75 | 4000* |
| | | Average | | -634 | +326 | |

*When output contained multiple consecutive significant blocks, they were merged for this analysis, resulting in blocks with total number SNPs in block greater than max window size

**Table 3. Difference in coordinates for significant blocks using sliding windows**

| Region Size | | Window | | # SNPs from start or end* | | Mean # SNPs in block |
|---|---|---|---|---|---|---|
| bp | SNPs | Size | Overlap | Mean | Max | |
| 400,000bp | 1086 | 2000 | 200 | 900 | 1799 | 2000 |
| 400,000bp | 1086 | 1000 | 200 | 400 | 799 | 1000 |
| 400,000bp | 1086 | 3000 | 200 | 1400 | 2799 | 3000 |
| 1,000,000bp | 2729 | 2000 | 200 | 900 | 1799 | 4000** |

*Theoretical mean and max distance

**When output contained multiple consecutive significant blocks, they were merged for this analysis, resulting in blocks with total number SNPs in block greater than max window size

**Supplementary Note 3. Identifying optimal window size to region size ratio for local substructure scan.** We noticed that the window size may be an important factor in the type I error and power of local substructure scans with *summix-local*. We assessed the optimal window size based on the size of the region to be detected. To this end, we performed a simulations to detect a 400kb region with 3% difference in IAM-like proportions compared to the remainder of the chromosome, varying the window size in the fast catchup algorithm. We varied the window size from 100kb to 1,500kb (0.25 to 3.75x the region size). We find the highest power and lowest type 1 error when the window size is 0.5 to 2 times the size of the region to be detected (Supplementary Fig. 3).

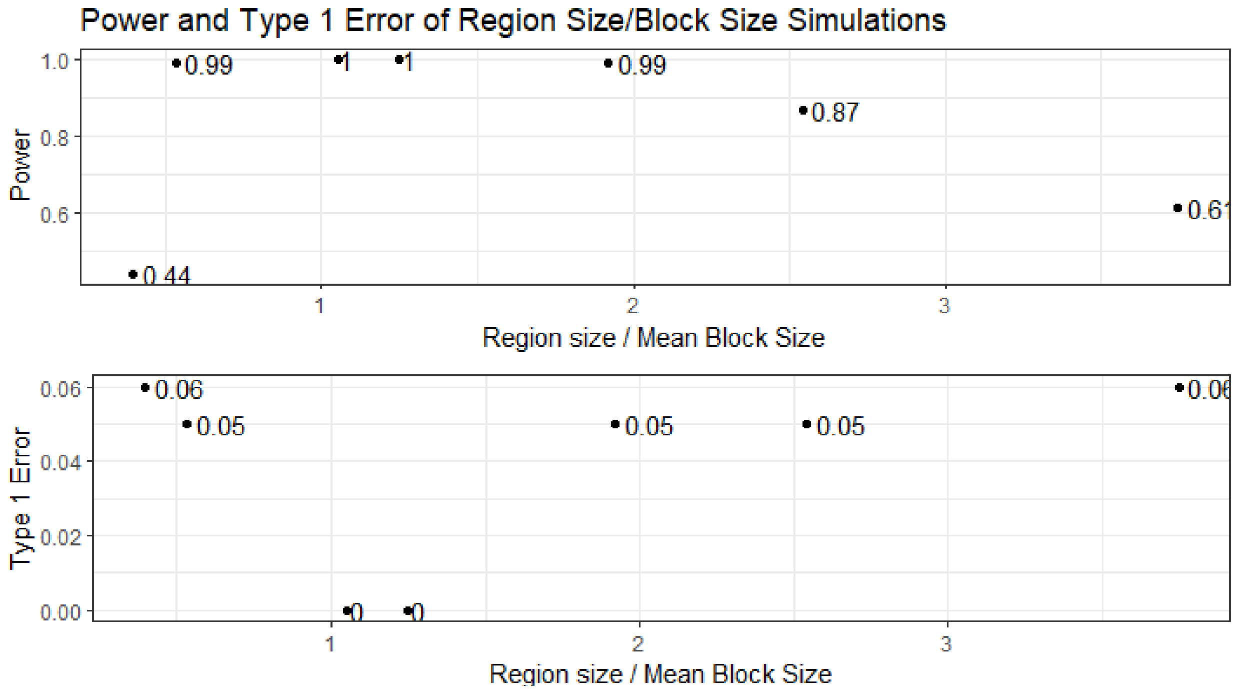


**Supplementary Fig. 3: Simulations show optimal window size is 0.5 to 2 times the size of the region to be detected.** Simulations were performed to determine the optimal window size to detect a 400Kb region with a 3% difference in substructure proportion. We find that the highest power and type I error is maintained when the window size is half to double the size of the region with different substructure proportions.

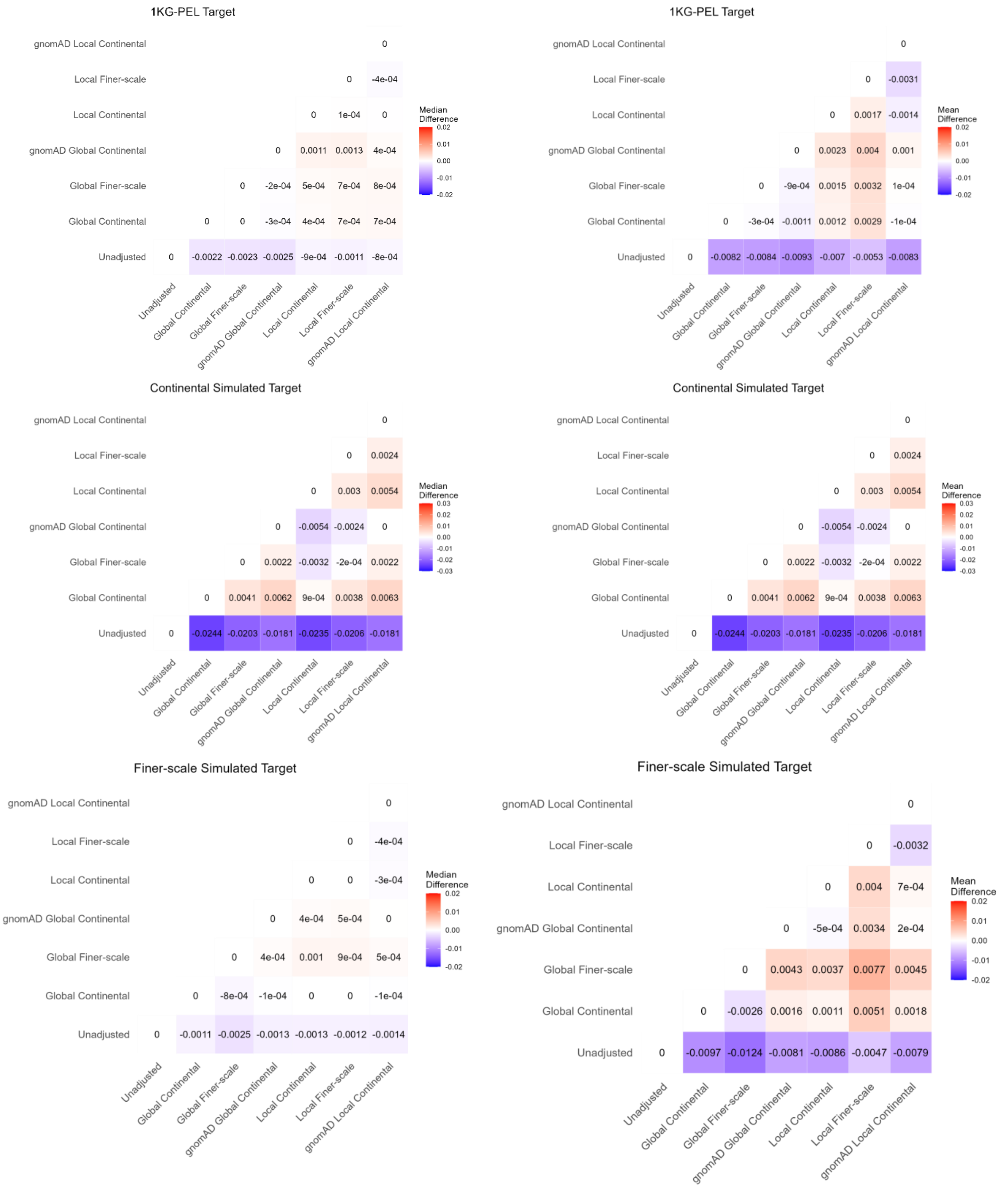


**Supplementary Fig. 4: Median and mean of absolute difference between observed and target AFs.** The median (left) and mean (right) absolute difference between the observed (unadjusted or adjusted) and the target AFs show that the improvement in the adjusted AFs is usually an order of magnitude or more than differences between the different types of adjustment.

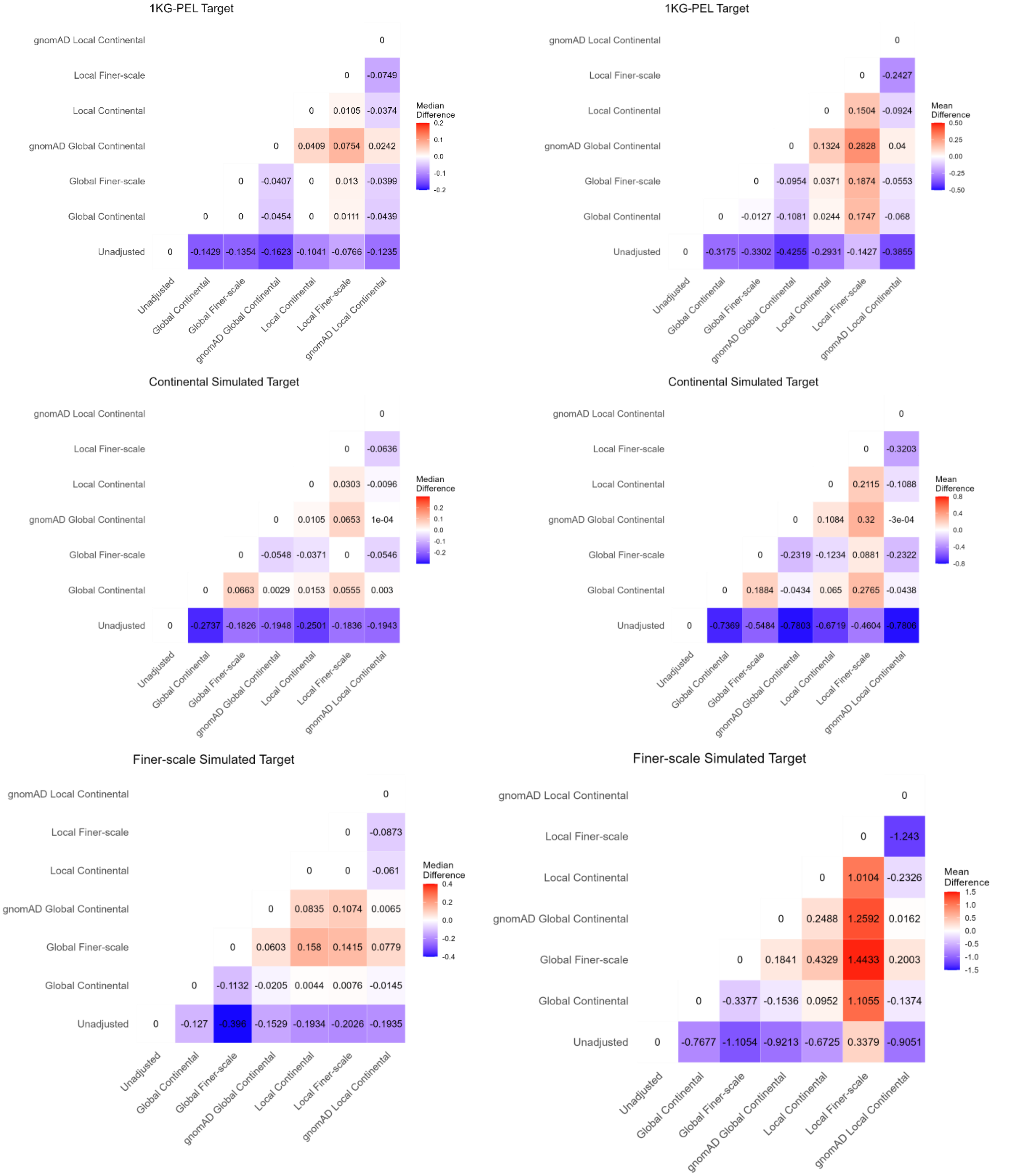


**Supplementary Fig. 5: Mean and median of relative difference between observed and target AFs.** The mean and median relative difference between the observed (unadjusted or adjusted) and the target AFs show that a large difference between the adjusted and unadjusted for all adjustments is an order of magnitude larger, indicating a large improvement. Notably, the local finer-scale adjustment for the finer-scale simulated target still contains large differences, likely due to confounding effects of the small reference sample size and small number of variants per window.

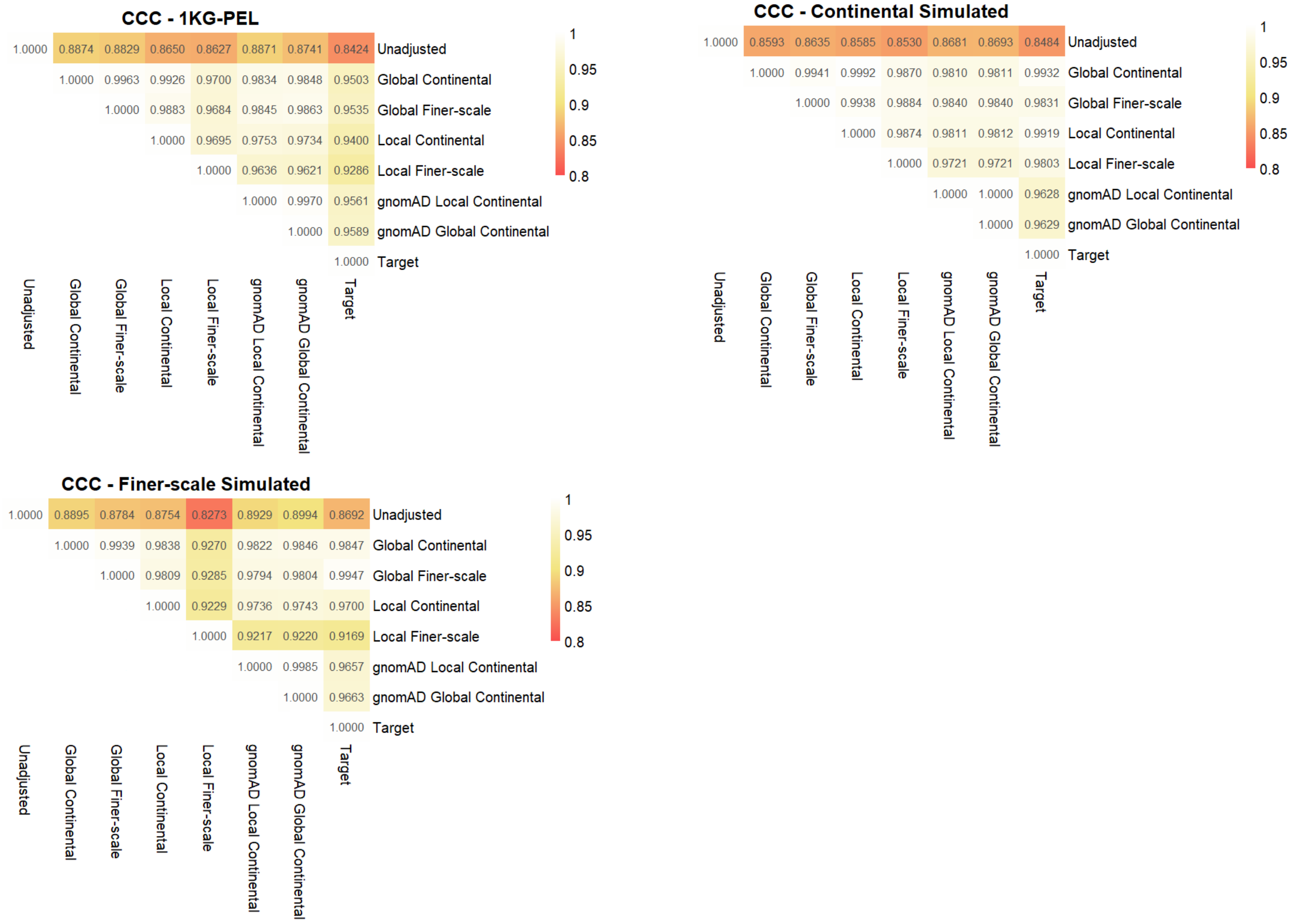


**Supplementary Fig. 6: All adjustments improve concordance between observed AFs and target AFs.** Lin's Concordance Correlation Coefficient (CCC) was estimated for each pair of target and observed (adjusted or unadjusted) AFs for the three adjustment scenarios (12,513 variants). All adjustments show improved (closer to 1.0) CCC with the target than the unadjusted AFs.

**Supplementary Note 4. Effective Sample Size Calculation**

*Summix2* estimates an effective sample size for the leave one reference group out *Summix* AF adjustment method as well as for the average AF adjustment method. Both effective sample size calculations are a weighted combination of the observed, $N_{obs}$, and reference sample sizes, $N_{ref,k}$. The first term in equation (1), the effective sample size calculation for the leave one out AF adjustment method, represents the information retained from the observed AF and transferred to the target sample while the second term is the information added from the reference data.

$$N_{eff,Leave\ Out\ l} = \min(\widehat{\pi_l}, \pi_{target,l}) * N_{obs} + \sum_{k \neq l}^{K} \pi_{target,k} * N_{ref,k} \quad (1)$$

The effective sample size estimation for the average AF adjustment method, equation (2), sums across K reference groups indicating additional information being gained by averaging over iteratively leaving out each reference group.

$$N_{eff,average} = \sum_{k=1}^{K} \min(\widehat{\pi_k}, \pi_{target,k}) * N_{obs} + \pi_{target,k} * N_{ref,k} \quad (2)$$

Examples of effective sample size calculations, reflecting the AF adjustment examples shown in Supplementary Figures 11-12, are in Table 4. More examples are in Supplementary Table 14.

**Table 4. Effective Sample Size Examples.** Effective sample sizes using equation (1) and equation (2) for AF adjustment scenarios in Supplementary Figures 10, 11, and 12.

| | Observed props | | | | | Target props | | | | | | | |
|---|---|---|---|---|---|---|---|---|---|---|---|---|---|
| AF Adjustment Scenario | AFR-like | EAS-like | EUR-like | IAM-like | SAS-like | AFR-like | EAS-like | EUR-like | IAM-like | SAS-like | $N_{eff,average}$ | $l$ | $N_{eff,Leave\ Out\ l}$ |
| AMR-like to PEL-like | .094 | 0.035 | 0.586 | 0.276 | 0.009 | 0.03 | 0.04 | 0.19 | 0.727 | 0.013 | 5900 | EUR-like | 2305 |
| | | | | | | | | | | | | IAM-like | 2896 |
| AFRAM-like to AFR-like | .84 | -- | .16 | -- | -- | 1 | -- | 0 | -- | -- | 8900 | AFR-like | 8000 |
| AFRAM-like to AFRAM-like | .5 | -- | .5 | -- | -- | .8 | -- | .2 | -- | -- | 7500 | AFR-like | 5100 |
| | | | | | | | | | | | | EUR-like | 2400 |

Sample size for simulated observed groups: $N_{obs} = 10{,}000$
Sample sizes for simulated reference groups: $N_{ref,k} = 500$

The effective sample size estimates are largest for the average AF adjustment method (Table 4). This is expected given the average AF adjustment method retains information from all reference groups, as opposed to the leave one reference group out, which retains only the information from the $l^{th}$ group in the observed AF. Of note, leaving out the reference group with the largest mixing proportion in the observed data will result in smaller effective sample sizes in most cases. A warning is produced by the *Summix2* AF adjustment function if either the effective sample size calculated with equation (1) or equation (2) is less than 50% of the sample size of the observed group.

**Supplementary Note 5. Calculating reference case AFs from Prostate Cancer Summary Statistics.**

We selected a prostate cancer GWAS that published odds ratio (OR) estimates and control AF for each variant. Using the case and control sample sizes ($N_{case}, N_{control}$) we calculated allele counts (AC) for the effect and non-effect alleles using $AC = AF * N$. We use the classic OR equation, equation (3), below to solve for the AC of the effect allele in cases, equation (4), as needed for our reference data:

$$OR = \frac{AC_{effect,case} * AC_{non-effect,control}}{AC_{effect,control} * AC_{non-effect,case}} \quad (3)$$

$$AC_{effect,case} = \frac{OR * AC_{effect,control}}{1 - AC_{effect,control} + OR * AC_{effect,control}} \quad (4)$$

**Supplementary Note 6. Updated allele frequency adjustment.**
In the Summix 2021 publication, users chose a reference group, $l$, to remove from the AF adjustment (equation (5)).

$$x = \frac{\pi_{target,l}}{\widehat{\pi_l}}\left(AF_{observed} - \sum_{k\neq l}\widehat{\pi_k}AF_{ref,k}\right) + \sum_{k\neq l}\pi_{target,k}AF_{ref,k} \quad (5)$$

$$AF_{adj} = \begin{cases} 0, & x < 0 \\ x, & 0 \leq x \leq 1 \\ 1, & x > 1 \end{cases}$$

Where $l$ is the index for the reference group left out and $\pi_{target,l}$ and $\widehat{\pi_l}$ refer to the target and observed group mixing proportions for $l$, respectively. $K$ is the reference group index, $\pi_{target,k}$ and $\widehat{\pi_k}$ refer to the target and observed group mixing proportions for $k$, $AF_{observed}$ is the AF vector for the observed group, $AF_{ref,k}$ is the AF vector for reference group $k$ , and $AF_{adj}$ is the adjusted AF.

In the AF adjustment examples shown within the 2021 publication, we chose $l$ to be the most common reference group in the observed summary data. In *Summix2*, we update the default AF adjustment method to iteratively exclude reference groups and average the adjusted AFs (equation (6)).

$$x^* = \frac{1}{K}\sum_{l=1}^{K}(x) \quad (6)$$

$$AF_{adj} = \begin{cases} removed, & x^* < \delta \\ 0, & \delta \leq x^* < 0 \\ x^*, & 0 \leq x^* \leq 1 \\ 1, & x^* > 1 \end{cases}$$

Where $x$ is from equation (5) and $\delta$ is the precision threshold for which adjusted AFs are removed from the dataset or rounded to 0.

In the development of *Summix2*, we wanted a default AF adjustment that did not require users to choose a reference group, $l$, to leave out. We explored a variety of new AF adjustment methods; each iterating through equation (5), leaving out a different reference group, $l$, per iteration and either averaging across all resulting adjusted AFs (equation (6)), weighting the resulting adjusted AFs by the $l$th reference group target proportion ($\pi_{target,l}$) (equation (7)), or weighting the resulting adjusted AFs by the $l$th reference group's sample size ($N_{ref,l}$) proportion (equation (8)).

$$x^* = \sum_{l=1}^{K}\pi_{target,l}(x) \quad (7)$$

$$x^* = \sum_{l=1}^{K}\frac{N_{ref,l}}{\sum_{l}^{K}N_{ref,l}}(\text{x}) \quad (8)$$

We assessed the performance of each of the AF adjustment methods in equations (6)-(8) across a variety of simulation scenarios. We found the average AF adjustment method (equation (6)) and the original Summix method of leaving out $l$ (equation (5)), given that $l$ is the reference group with the largest proportion in the observed sample, to be the most robust across all simulation scenarios (Supplementary Figures 7-9). In particular, the AF adjustment from AMR-like to IAM-like using finer-scale reference groups performs best when using the average AF adjustment method (Supplementary Figure 9). This is likely due to the mixing proportion of each reference group only representing a small proportion of the observed AF (i.e., the largest $\widehat{\pi_k}$ is seen for $\hat{\pi}_{Maya-like} = 0.17$). Full simulated observed and target group proportions in Supplementary Table 11.

We then compared the performance of the original AF adjustment (equation (5)) to the average AF adjustment (default in *Summix2*, equation (6)). We simulated observed AMR-like AFs and adjusted to simulated target PEL-like AFs and assessed the absolute difference between the target AF and the AFs adjusted using the original *Summix* adjusted AF method when leaving out IAM-like, EUR-like, EAS-like, AFR-like, and SAS-like, and the average AF adjustment method (Supplementary Figure 10). We find that when leaving out IAM-like, the most common reference group present in the target group summary data, there is higher absolute difference compared to either leaving out each of the other reference groups or using the average adjusted AF. This is likely because IAM-like is not the most common reference in the observed data and has the largest proportion increase from the observed to target groups ($\hat{\pi}_{IAM-like}$= 0.276 to $\pi_{target,IAM-like}$= 0.727). We find that the average AF adjustment performs similarly to when the reference group with the largest observed mixing proportion (EUR, $\hat{\pi}_{EUR-like}$= 0.586) is left out. Similarly, AF adjustments from an AFR/AFRAM-like observed group ($\hat{\pi}_{AFR-like} = 0.8, \hat{\pi}_{EUR-like} = 0.2$) to an AFR-like target group ($\pi_{target,AFR-like} = 1$), (Supplementary Figure 11), perform best when using the average AF adjustment and leaving out the most common reference group in the observed sample (i.e., AFR-like). When considering an AF adjustment from an AFR/AFRAM-like observed group ($\hat{\pi}_{AFR-like} = 0.5, \hat{\pi}_{EUR-like} = 0.5$) to an AFR/AFRAM-like target group ($\pi_{target,AFR-like} = .8, \pi_{target,EUR-like} = .2$) (Supplementary Figure 12), there is not a reference group with the largest proportion in the observed sample. Here, the average Summix AF adjustment performs the best.

Overall, we found that the average AF adjustment method and the original AF adjustment method (leaving out the largest reference group in the observed data) performed best across all simulation scenarios. The average AF adjustment method is the default as it does not require user expertise, although both adjustments are options in the *Summix2* AF adjustment function. We especially recommend using the default average AF adjustment when: (1) there is a substantial change in mixing proportions for the left out group (especially increasing from observed to target, e.g, Supplementary Figures 9 and 12) and (2) using finer-scale reference groups (Supplementary Figure 9).

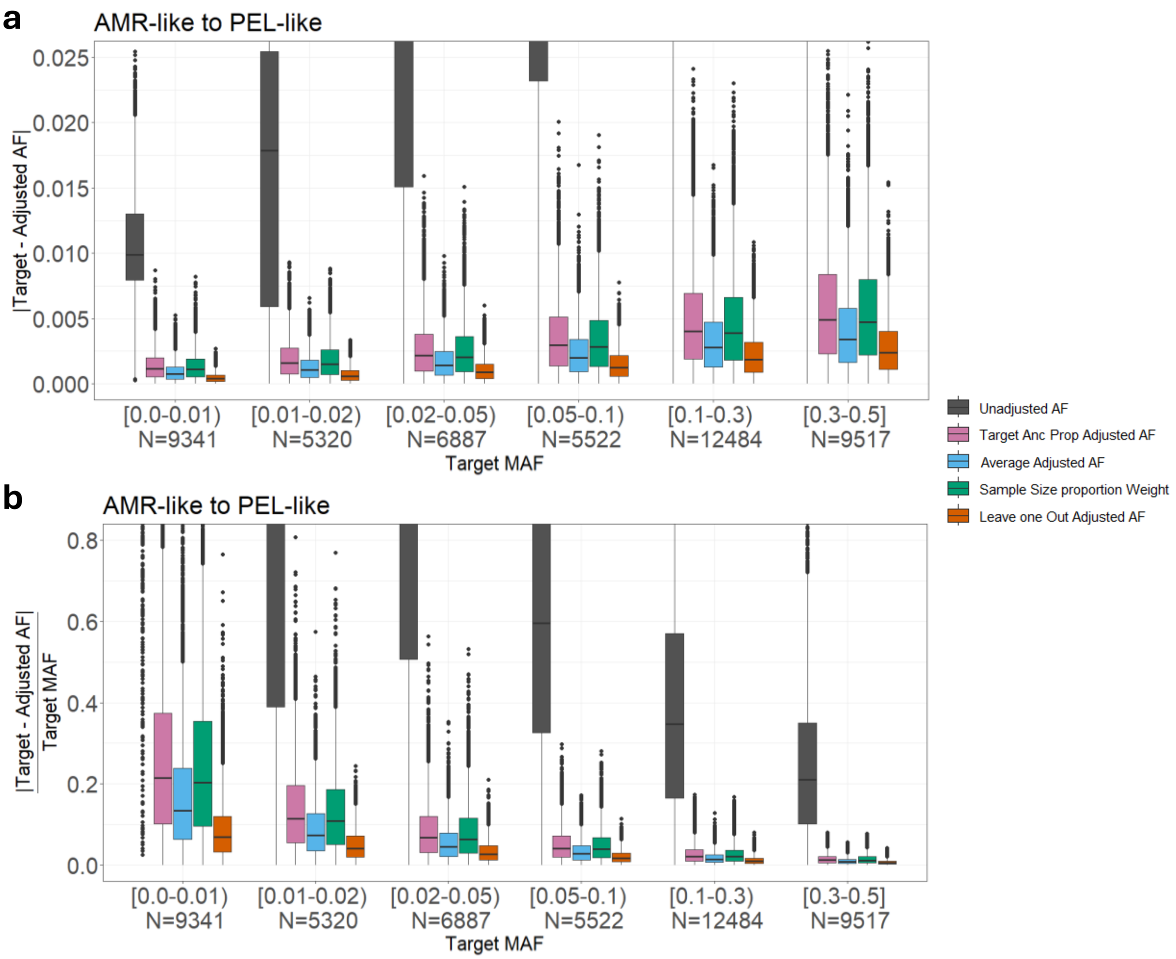


**Supplementary Fig. 7. Data harmonization for AMR-like samples with continental substructure.** Absolute difference (**a**), and relative difference (**b**), after adjusting AMR-like (EUR-like=.586, IAM-like=.276, AFR-like=.094, EAS-like=.035, SAS-like=.009) to PEL-like (EUR-like=.19, IAM-like=.727, AFR-like=.03, EAS-like=.04, SAS-like=.013) comparing unadjusted (grey), Summix adjusted with target mixing proportions (pink), *Summix2* average adjusted (light blue), Summix adjusted with reference sample size proportions (green), and Summix adjusted leaving out EUR-like (orange).

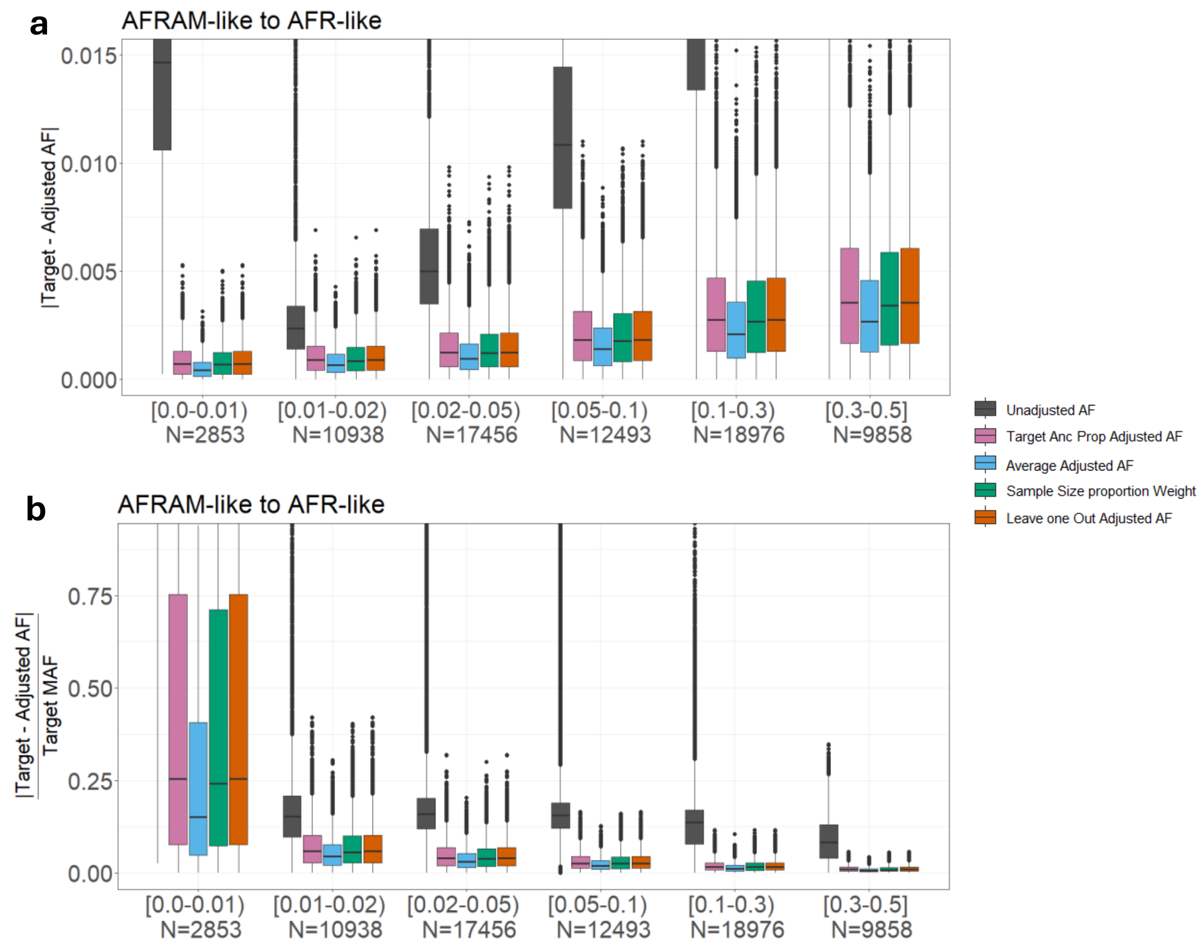


**Supplementary Fig. 8. Data harmonization for AFRAM-like samples with continental substructure.** Absolute difference (**a**), and relative difference (**b**), after adjusting AFRAM-like (AFR-like=.84, EUR-like=.16) to AFR-like (AFR-like=1) comparing unadjusted (grey), Summix adjusted with target mixing proportions (pink), *Summix2* average adjusted (light blue), Summix adjusted with reference sample size proportions (green), and Summix adjusted leaving out AFR-like (orange).

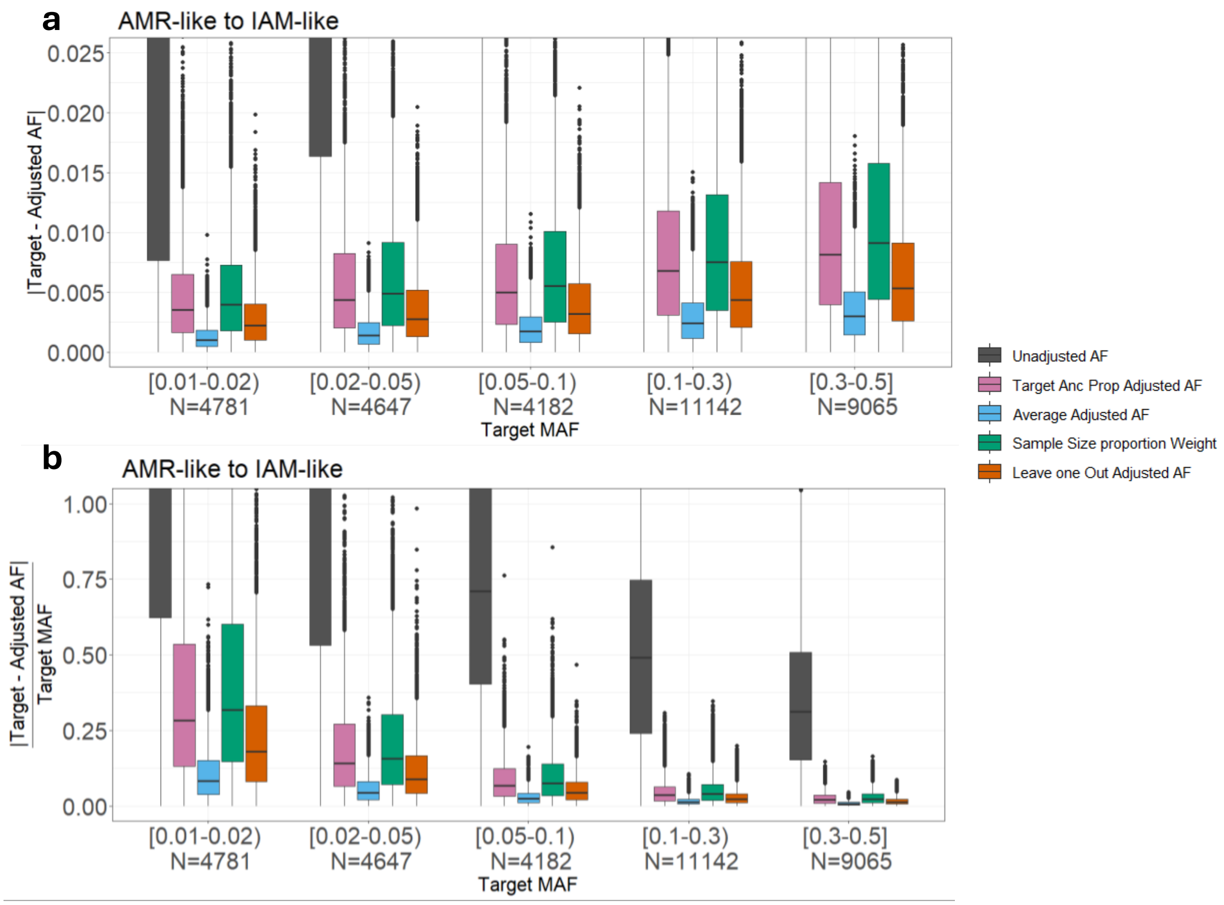


**Supplementary Fig. 9. Data harmonization for AMR-like samples with finer-scale substructure.** Absolute difference (**a**), and relative difference (**b**), after adjusting AMR-like to IAM-like (simulated finer-scale proportions in Supplementary Table 11) comparing Unadjusted (grey), Summix adjusted with target mixing proportions (pink), *Summix2* average adjusted (light blue), Summix adjusted with reference sample size proportions (green), and Summix adjusted leaving out Maya-like (orange).

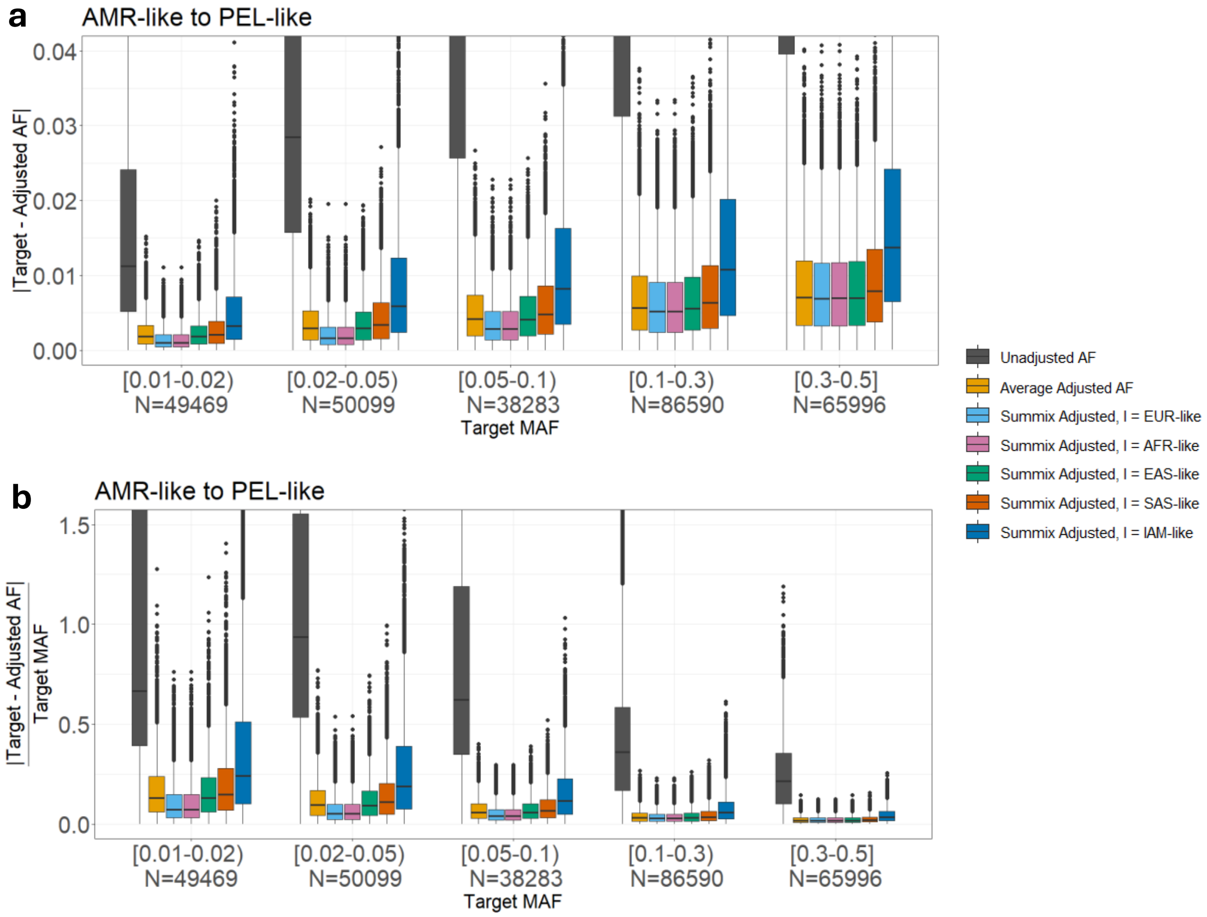


**Supplementary Fig. 10. Comparing *Summix2* data harmonization for average and leave one reference group out for AMR-like sample with continental substructure.** Absolute difference (**a**), and relative difference (**b**), after adjusting AMR-like (EUR-like=.586, IAM-like=.276, AFR-like=.094, EAS-like=.035, SAS-like=.009) to PEL-like (EUR-like=.19, IAM-like=.727, AFR-like=.03, EAS-like=.04, SAS-like=.013) comparing unadjusted (grey), *Summix2* average adjusted (light orange), Summix adjusted leaving out EUR-like (light blue), Summix adjusted leaving out AFR-like (pink), mix adjusted leaving out EAS-like (green), Summix adjusted leaving out SAS-like (dark orange), and Summix adjusted leaving out IAM-like (dark blue).

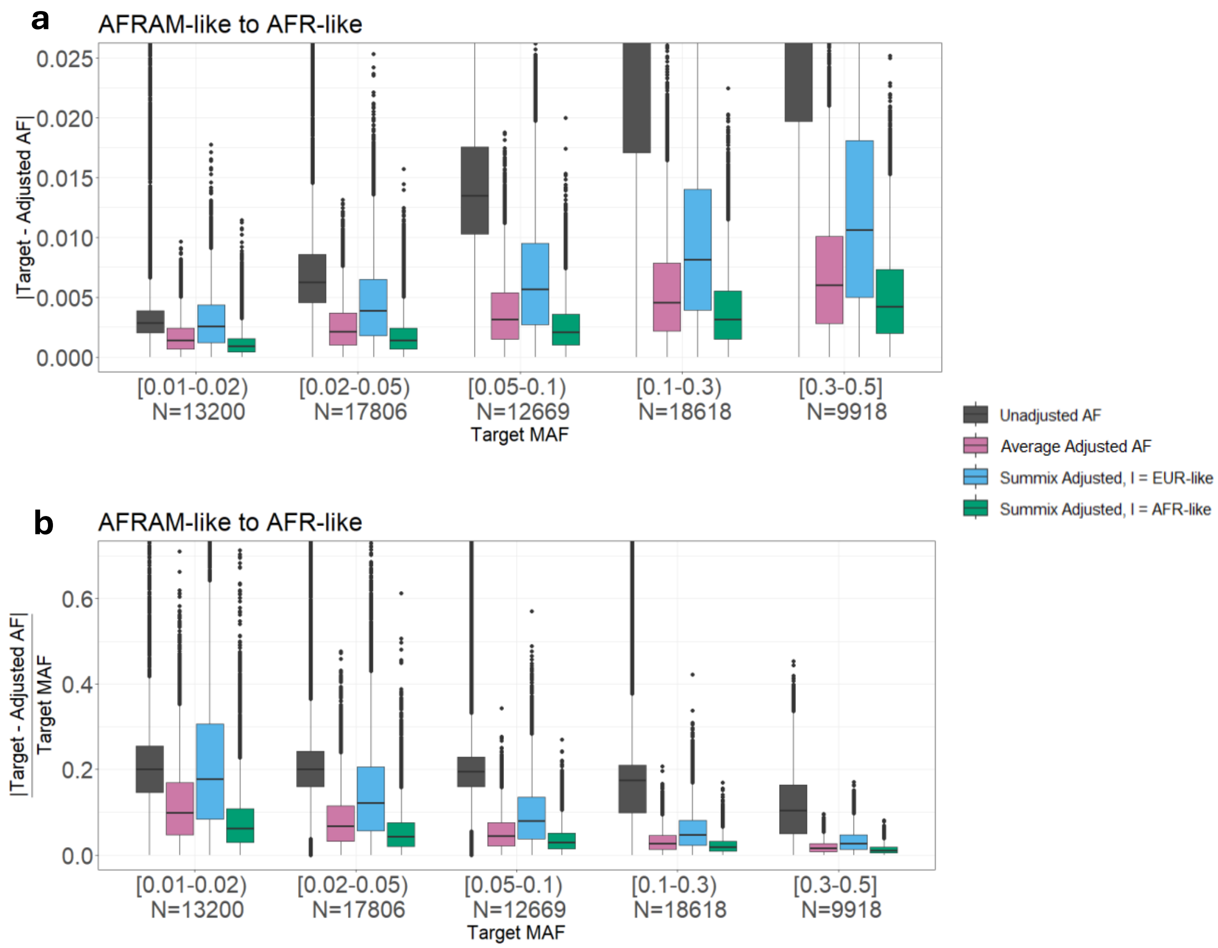


**Supplementary Fig. 11. Comparing *Summix2* data harmonization for average and leave one reference group out for AFRAM-like sample with continental substructure.** Absolute difference (**a**), and relative difference (**b**), after adjusting AFRAM-like (AFR-like=.8, EUR-like=.2) to AFR-like (AFR-like=1) comparing Unadjusted (grey), *Summix2* average adjusted (pink), Summix adjusted leaving out EUR-like (light blue), and Summix adjusted leaving out AFR-like (green).

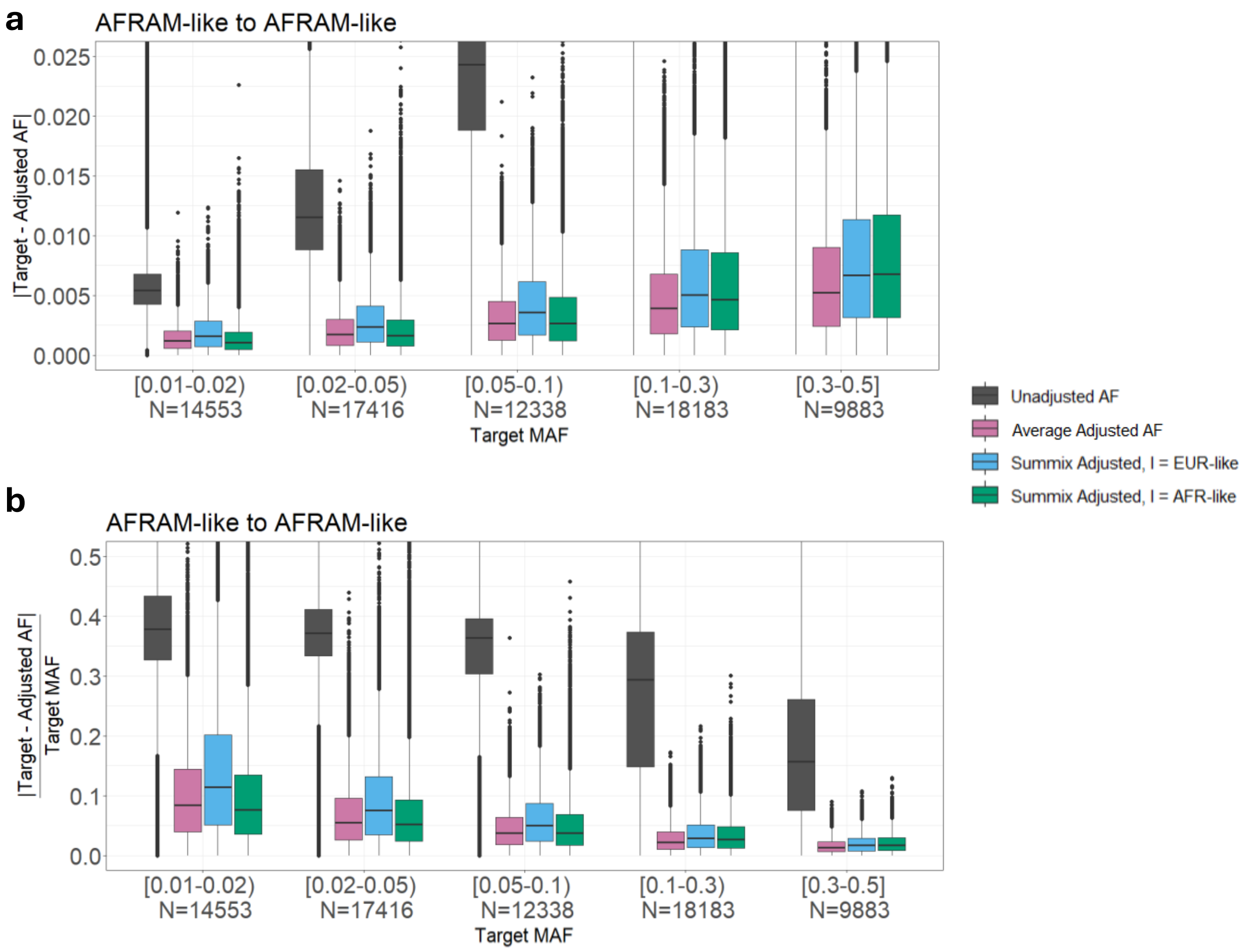


**Supplementary Fig. 12. Comparing *Summix2* data harmonization for average and leave one reference group out for AFRAM-like sample with 50%-50% continental substructure.** Absolute difference (**a**), and relative difference (**b**), after adjusting AFRAM-like (AFR-like=.5, EUR-like=.5) to AFRAM-like (AFR-like=.8, EUR-like=.2) comparing Unadjusted (grey), *Summix2* average adjusted (pink), Summix adjusted leaving out EUR-like (light blue), and Summix adjusted leaving out AFR-like (green).

**Supplementary Note 7. Removing reference groups with small global proportion estimates from local substructure estimation.**

Through empirical, we found that using all five continental references (AFR, EUR, EAS, IAM, SAS) when the substructure was not present on average across the genome resulted in high variability in local substructure estimates elevating the false positive rate for the reference groups with small global proportions. Removing reference groups contributing <2% to the global substructure eliminated these false positives. Therefore, by default in *Summix_local*, we check for global contribution of each reference group and remove those with <2%.
